# Mapping the Functional Landscape of the Receptor Binding Domain of T7 Bacteriophage by Deep Mutational Scanning

**DOI:** 10.1101/2020.07.28.225284

**Authors:** Phil Huss, Anthony Meger, Megan Leander, Kyle Nishikawa, Srivatsan Raman

## Abstract

The interaction between a bacteriophage and its host is mediated by the phage’s receptor binding protein (RBP). Despite its fundamental role in governing phage activity and host range, the molecular rules of RBP function remain a mystery. Here, we systematically dissect the functional role of every residue in the tip domain of T7 phage RBP using a novel phage genome engineering method called ORACLE (**<u>O</u>**ptimized **<u>R</u>**ecombination, **<u>Ac</u>**cumulation and **<u>L</u>**ibrary **<u>E</u>**xpression). ORACLE is a high-throughput, locus-specific, sequence-programmable method to create a large, unbiased library of phage variants at a targeted gene locus. Using ORACLE, we generated all single amino acid substitutions at every site (1660 variants) of the tip domain to quantify the functional role of all variants on multiple bacterial hosts. This rich dataset allowed us to cross compare functional profiles of each host to precisely identify regions of functional importance, many which were previously unknown. Host-specific substitution patterns displayed differences in site specificity and physicochemical properties of mutations indicating exquisite adaptation to individual hosts. Comparison of enriched variants across hosts also revealed a tradeoff between activity and host range. We discovered gain-of-function variants effective against resistant hosts and host-constricting variants that selectively eliminated certain hosts. We demonstrate therapeutic utility against uropathogenic *E. coli* by engineering a highly active T7 variants to avert emergence of spontaneous resistance of the pathogen. Our approach presents a generalized framework for systematic and comprehensive characterization of sequence-function relationships in phages on an unprecedented scale.

## Introduction

Bacteriophages (or “phages”) shape microbial ecosystems by infecting and killing targeted bacterial species. As a result, they are promising tools for treatment of antibiotic resistant bacterial infections and microbiome manipulation [1]–[12]. Interaction of phages with their bacterial receptors is a key determinant of their host range and virulence [13]–[15]. This interaction is primarily mediated by the receptor binding proteins (RBPs) of the phage [16]. RBPs enable phages to adsorb to diverse cell surface molecules including proteins, polysaccharides, lipopolysaccharides and carbohydrate-binding moieties. Phages exhibit high functional plasticity through genetic alterations to RBPs and by natural and laboratory-guided evolution which can modulate activity and host range to different hosts and environments [4], [17]–[26]. In essence, survivability of a phage is intimately linked to the adaptability of its RBP. The challenge now is to understand the molecular code of RBPs in sufficient depth to enable manipulation of host range and virulence predictably. We sought to do so by combining deep mutational scanning of the RBP with powerful selections on multiple hosts.

Although RBPs remain the focus of many mechanistic, structural and evolutionary studies and are a prime target for engineering, we currently lack a systematic and comprehensive understanding of how RBP mutations influence phage activity and host range. Though insightful, directed evolution enriches only a small group of ‘winners’ which makes it difficult to glean a comprehensive mutational landscape of the RBP [22]. Random mutagenesis-based screens generate multi-mutant variants whose individual effects cannot be easily deconvolved [19], [25]. Other approaches including swapping homologous RBPs lead to gain of function, however the underlying molecular determinants of function can be difficult to explain [17], [18], [21], [26]. In summary, despite the extraordinary functional potential of phage RBPs, how systematic changes to their sequence shape the overall functional landscape of a phage remains unknown.

Here we carried out deep mutational scanning (DMS), a high-throughput experimental technique, of the tip domain of the T7 phage RBP (tail fiber) to uncover molecular determinants of activity and host range. The tip domain is the distal region of the tail fiber that makes primary contact with the host receptor [27]–[29]. We developed ORACLE (**<u>O</u>**ptimized **<u>R</u>**ecombination, **<u>Ac</u>**cumulation and **<u>L</u>**ibrary **<u>E</u>**xpression), a high-throughput, locus-specific, phage genome engineering method to create a large, unbiased library of phage variants at a targeted gene locus. Using ORACLE, we systematically and comprehensively mutated the tip domain by making all single amino acid substitutions at every site (1660 variants) and quantified the functional role of all variants on multiple bacterial hosts. We generated high resolution functional maps delineating regions concentrated with function-enhancing substitutions and host-specific substitutional patterns, many of which were previously unknown. We discovered T7 variants with far greater virulence than wildtype T7, demonstrating that even natural phages well adapted to a host can be engineered for higher efficacy. However, many variants highly adapted to one host performed poorly on others, underscoring a tradeoff between activity and host range. This functional screening highlights ideal regions of the tip domain for engineering host range. Furthermore, we demonstrated the functional potential of RBPs by discovering gain-of-function variants against resistant hosts and host-constriction variants that selectively eliminate specific hosts. To demonstrate the therapeutic value of ORACLE, we engineered T7 variants that avert emergence of spontaneous resistance in pathogenic *E. coli* causing urinary tract infections.

Our study explains the molecular drivers of adaptability of the tip domain and identifies key functional regions determining activity and host range. ORACLE provides a generalized framework to describe sequence-function relationships in phages to demystify the molecular basis of phages, the most abundant life form on earth.

## Results

### Creating an unbiased library of phage variants using ORACLE

ORACLE is a high-throughput precision phage genome engineering technology designed to create a large, unbiased library of phage variants to investigate sequence-function relationships in phages. ORACLE overcomes three major hurdles. First, phage variants are created during the natural infection cycle of the phage which eliminates a common bottleneck from transforming DNA libraries. By recombining a donor cassette containing prespecified variants to a targeted site on the phage genome, ORACLE allows sequence programmability and generalizability to virtually any phage. Second, ORACLE minimizes library bias that can rapidly arise due to fitness advantage or deficiency of any variant on the propagating host that may then be amplified due to exponential phage growth. Minimizing bias is critical because variants that perform poorly on a propagating host but well on targeted hosts may disappear during propagation. Third, ORACLE prevents extreme abundance of wildtype over variants, which allows for resolving and scoring even small functional differences between variants. The development of this technology was necessary to overcome challenges with existing engineering approaches for creating a large, unbiased phage library. Direct transformation of phage libraries, while ideal for creating one or small groups of synthetic phages, will not work because phage genomes are typically too large for library transformation or are reliant on highly transformable hosts [17], [30]–[32]. Homologous recombination has low, variable recombination rates and high levels of wildtype phage are retained, which mask library members [25], [33]. Libraries of lysogenic phages could potentially be made using conventional bacterial genome engineering tools as the phage integrates into the host genome. However, this approach is not applicable to obligate lytic phages. Our desire to develop ORACLE for obligate lytic phages is motivated by their mandated use for phage therapy. Any phage, including lysogenic phages, with a sequenced genome and a plasmid-transformable propagation host should be amenable to ORACLE.

ORACLE is carried out in four steps: (a) making acceptor phage (b) inserting gene variants through recombination (c) accumulating recombined phages (d) expressing the library for selection (Figure 1A). An ‘acceptor phage’ is a synthetic phage genome where the gene of interest (i.e., tail fiber) is replaced with a fixed sequence flanked by Cre recombinase sites to serve as a landing site for inserting variants (Figure S1). T7 acceptor phages lacking a wildtype tail fiber gene cannot plaque on *E. coli* and do not spontaneously re-acquire the tail fiber during propagation (Figure 1B and Figure S2A). Furthermore, the T7 acceptor phages have no plaquing deficiency relative to wildtype when the tail fiber gene is provided from a donor plasmid (Figure S2A). Thus, the tail fiber gene is decoupled from the rest of the phage genome for interrogation of function. Next, phage variants are generated within the host during the infection cycle by **<u>O</u>**ptimized **<u>R</u>**ecombination by inserting tail fiber variants from a donor plasmid into the landing sites in the acceptor phage. To minimize biasing of variants during propagation, a helper plasmid constitutively provides the wildtype tail fiber *in trans* such that all progeny phages can amplify regardless of the fitness of any variant. At this stage, we typically have ∼1 recombined phage among 1000 acceptor phages (Figure 1C). To enrich recombined phages in this pool, we passage all progeny phages on *E. coli* expressing Cas9 and a gRNA targeting the fixed sequence flanked by recombinase sites we introduced into the acceptor phage. As a result, only unrecombined phages will be inhibited while recombined phages with tail fiber variants are **<u>Ac</u>**cumulated. The Cas9-gRNA system has no effect on plaquing of untargeted phages (Figure 2B and S2B-D). Recombined phages were highly enriched by over 1000-fold in the phage population when an optimized gRNA (Figure S2A) targeting the fixed sequence was used, whereas a randomized control gRNA yielded no enrichment of recombined phages (Figure 1D and S2E-F). In the final step, phages which were thus far propagated on hosts complementing wildtype tail fiber on a helper plasmid, are propagated on *E. coli* lacking the helper plasmid for **<u>L</u>**ibrary **<u>E</u>**xpression of the variant tail fiber on the recombined phage. The distribution of the library of tail fiber variants integrated on the phage genome after ORACLE was mildly skewed towards more abundant members but remained generally evenly distributed and comparable to the distribution of variants in the input donor plasmid library, retaining 99.8% coverage (Figure 1E, Table S1 and S3). Comparison of variant libraries with and without DNAse treatment was well correlated (R=0.994), indicating no unencapsidated phage genomes influenced library distribution (Figure S3A). In summary, ORACLE is a generalizable tool for creating large, unbiased variant libraries of obligate lytic phages. These phage variants, including those that have a fitness deficiency on the host used to create the library, can all be characterized in a single selection experiment by deep sequencing phage populations before and after selection in a host. Compared to traditional plaque assays this represents increased throughput by nearly 3-4 orders of magnitude.

**Figure 1.**
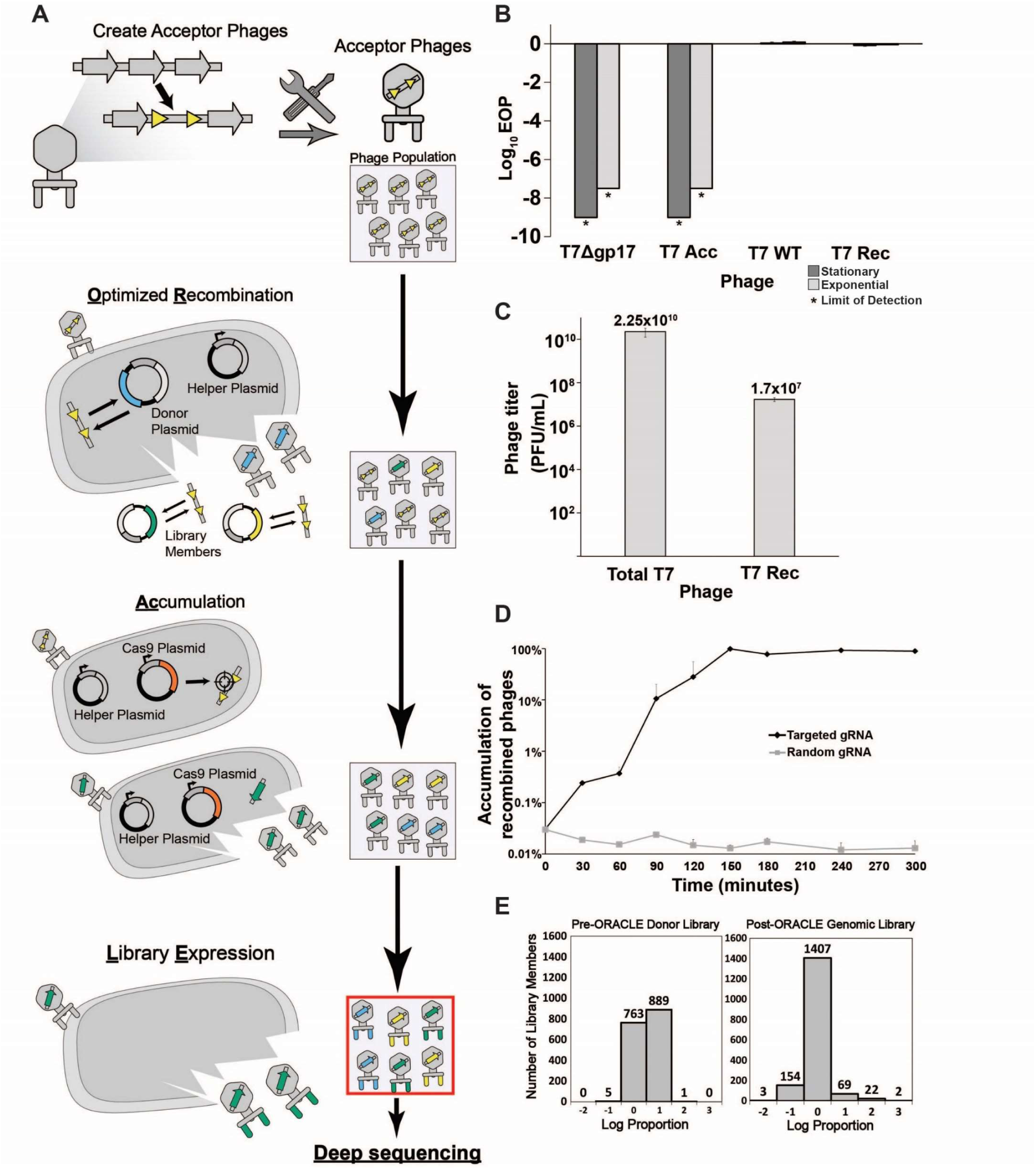
ORACLE workflow for creating phage variant libraries. **(A)** Schematic illustration of the four steps of ORACLE: creation of acceptor phage, inserting gene variants (**<u>O</u>**ptimized **<u>R</u>**ecombination), enriching recombined phages (**<u>A</u>**ccumulation) and expressing library for selection (**<u>L</u>**ibrary **<u>E</u>**xpression). Color notations are as follows: yellow triangles – Cre recombinase sites, blue, green and yellow colored segments – gene variants, orange segment – Cas9. **(B)** Ability of different versions of T7 to infect *E. coli* 10G in exponential (dark gray bar) and stationary (light gray bar) phases by EOP using exponential 10G with *gp17* tail fiber helper plasmid as reference host. T7 without tail fiber (T7Δgp17) and T7 without tail fiber (T7Δgp17) plaque poorly, but wildtype T7 (T7 WT), and T7 with *gp17* recombined into the acceptor locus (T7 Rec) plaque efficiently. **(C)** Concentration of total (Total T7) and recombined (T7 Rec) phages after a single passage on host containing Cre recombinase system. Recombination rate is estimated to be ∼7.19×10^-4^. **(D)** Percentage of recombined phages in total phages when using gRNA targeting fixed sequence at acceptor site T7 Acc (Targeted) or randomized gRNA (Random). **(E)** Histogram of abundance of variants in the input plasmid library (left) and on the phage genome after ORACLE (right) binned using log proportion centered on equal representation. All data represented as mean ± SD of biological triplicate. See also Figure S1 and S2 and Table S3.

**Figure 2.**
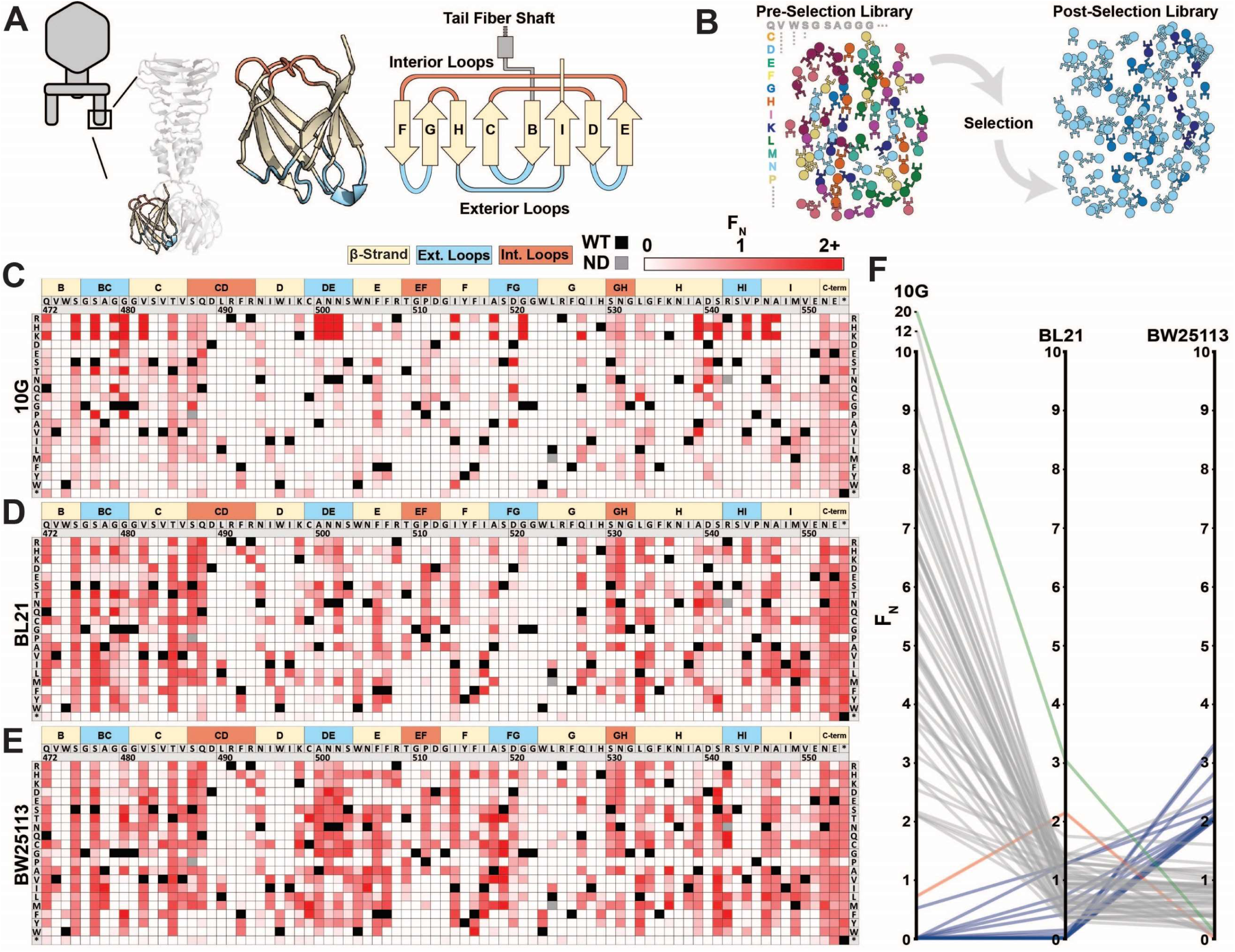
Deep mutational scanning of tip domain shows phage adaptation at molecular resolution. **(A)** Crystal structure and secondary structure topology of the tip domain color coded as interior loops (red), β-sheets (beige) and exterior loops (blue) **(B)** Functional analysis of variants by comparing their abundances pre-and post-selection on a host. **(C-E)** Heat maps showing normalized functional scores (F_N_) of all substitutions (red gradient) and wildtype amino acid (solid black) at every position for *E. coli* 10G (**C**), BL21 (**D**) and BW25113 (**E**). Residue numbering (based on PDB 4A0T), wildtype amino acid and secondary structure topology are shown above left to right, substitutions listed top to bottom. **(F)** Parallel plot showing F_N_ for enriched (F_N_≥2) variants on 10G, BL21, and BW25113. Coloring indicates enrichment only on 10G (grey), only on BL21 (red), only on BW25113 (blue) enriched on 10G and BL21 (green). Connecting lines indicate F_N_ of the same variant on other hosts. See also Figure S3 and Figure S4 and Table S2 and S3.

### Deep mutational scanning of tip domain shows phage adaptation at molecular resolution

Deep mutational scanning (DMS) is a high-throughput experimental technique to characterize sequence-function relationship through large-scale mutagenesis coupled to selection and deep sequencing. The scale and depth of DMS is used to reveal sites critical for activity, host specificity, and stability in a protein. DMS has been employed to study many proteins including enzymes, transcription factors, signaling domains, and viral surface proteins [34]–[37].

Bacteriophage T7 is a podovirus that infects *E. coli*. T7 has a short non-contractile tail made up of three proteins including the tail fiber encoded by *gp17*. Each of the six tail fibers is a homotrimer composed of a relatively rigid shaft ending with a β-sandwich tip domain connected by a short loop [38]. The tip domain is likely the very first region of the tail fiber to interact with host lipopolysaccharide (LPS) and position the phage for successful, irreversible binding with the host [27]–[29]. The tip domain is a major determinant of host range and activity and is often naturally exchanged between phages to readily adapt to new hosts [23], [39], [40]. Even single amino acid substitutions to this domain are sufficient to alter host range between *E. coli* strains [41]. Due to its critical functional role, we chose the tip domain to comprehensively characterize phage activity and host range by DMS.

We generated a library of 1660 single mutation variants of the tip domain, pre-specified as chip-based oligonucleotides, where all nineteen non-synonymous and one nonsense substitution were made at each codon spanning residue positions 472-554 (Figure 2A, residue numbering based on PDB 4A0T). Using ORACLE, the library was inserted into T7 to generate variants to be selected and deep sequenced (Figure 2B) on three laboratory *E. coli* hosts: B strain derivative BL21, K-12 derivative BW25113 and DH10B derivative 10G. Each variant was given a functional score, F, based on the ratio of their relative abundance before and after selection, which was then normalized to wildtype to yield F_N_ (Figure 2C-E, Table S2 and S3, see methods). Selection on each host gave excellent correlation across biological triplicates (Figure S3B-D). To validate the functional relevance of the screen, we hypothesized that the flexible C-terminal end (residue positions 552-554 and a three-residue extension if the stop codon is substituted) is unlikely to have any structural or host recognition role. As expected, these positions broadly tolerated nearly all substitutions across all three hosts indicating that the functional scores likely reflect true biological effects (Figure 2C-E, Table S3).

We compared the activities of phage variants across hosts to assess their fitness and evolutionary adaptation to each host. Between the three hosts, T7 variants appeared most and least adapted to BW25113 and 10G, respectively, as evidenced by the fraction of depleted variants (F_N_<0.1) after selection on each host (10G:0.66, BL21:0.59, and BW25113:0.51) (Figure S4A-C). Furthermore, wildtype T7 fared relatively poorly on 10G (F=0.77), indicating a fitness impediment, but performed well on BL21 (F=2.92) and BW25113 (F=2.26) (Table S2). The fitness impediment gave many more variants competitive advantage resulting in greater enrichment (F_N_≥2) over wildtype on 10G (48 variants) compared to BL21 (2 variants) and BW25113 (16 variants) (Figure S4A-C). In fact, the best performing variants on 10G were 10 times more enriched than wildtype suggesting substantially higher activity (Figure 2F and S4D). Examining enriched variants on each host (F_N_≥2) provides compelling evidence of the tradeoff between activity and host range (Figure 2F, S4E and Table S3). The top ranked variants on each host were remarkably distinct from those on other hosts (except G479Q shared between 10G and BL21). Variants enriched on one host scored poorly on others (Figure 2F), but variants with intermediate scores performed well on all hosts (Table S3). Thus, specialization toward a host comes at the cost of sacrificing breadth, mirroring observations made of natural phage populations [42].

We investigated the physicochemical properties and topological preferences of substitutions after selection on each host (Figure S4F-H). On 10G, there was pronounced enrichment of larger and more hydrophilic amino acids and depletion of hydrophobic amino acids (all p<0.001 Figure S4F-G), which is visually striking on the heatmap (see R, K and H on Figure 2C). In contrast, no strong enrichment or depletion was observed on BL21 (Figure S4F-H). This is consistent with our earlier observation that wildtype T7 is generally well adapted to BL21 since it had the fewest variants outperforming wildtype. We reasoned that since BL21 has historically been used to propagate T7, it may have already adapted well to this host over time. On BW25113, hydrophobic residues were modestly enriched (p<0.036 Figure S4H), a trend opposite to 10G. This provides a molecular explanation as to why high scoring substitutions on one host fare poorly on others (Fig. 2F). We mapped positions of enriched substitutions (F_N_≥2) on each host onto the structure to determine topological preferences (Figure S4E). These fall predominantly on four exterior loops (BC, DE, FG, and HI), the adjoining region (β-strand H) close to exterior loop HI, and less frequently on the ‘side’ of the tip domain. This suggests directionality to phage-bacterial interactions and orientational bias of the tip domain with respect to the bacterial surface. Directionality and orientational bias is particularly valuable information since no high-resolution structure of this phage bound to receptor exists.

Several key lessons emerged from these host screens. First, single amino acid substitutions alone can generate broad functional diversity, highlighting the evolutionary adaptability of the RBP. Second, T7 can be optimized and activity can be increased, even on hosts that T7 is already considered to grow well on. Third, enrichment patterns on each host follow broad trends but have nuance at each position.

### Comparison across hosts reveals regions of functional importance

Next, we sought to elucidate features of each residue unique to each host or common across all hosts. There were over 30 residues with contrasting substitution patterns between different hosts, revealing fascinating features of receptor recognition for T7 (Table S3). Here we focus on five of these residues, N501, R542, G479, D540, and D520, which showed starkly contrasting patterns of selection (Figure 3A). N501 and R542 are located on exterior loops oriented away from the phage and toward the receptor (Figure 3C). In fact, R542 forms a literal ‘hook’ to interact with the receptor. On 10G and BL21, only positively charged residues (R, K and H) were tolerated at residues 501 and 542, while in contrast many more substitutions were tolerated at both residues on BW25113. One such substitution, R542Q, is the best performing variant on BW25113 (F_N_ = 3.31) but is conspicuously depleted on 10G and BL21 suggesting that even subtle molecular disparities can lead to large biases in activity. The substitution profiles of G479 and D540 are loosely the inverse of N501 and R542 as many substitutions are tolerated on BL21 and 10G, but very few are tolerated on BW25113 (Figure 3A). We hypothesize that D540 is critical for host-recognition on BW25113. Since D540, a receptor-facing position on an exterior loop, is only 6 Å from G479, it is likely that any substitution at G479 may sterically hinder D540, resulting in the noted depletion of G479 substitutions on BW25113. This hypothesis is further supported by enrichment of adjacent S541D on BW25113 (F_N_ = 2.82, the third highest scoring substitution), while this substitution is depleted on 10G and BL21 (Table S3). D520 displays a third variation in substitution patterns where substitutions are generally tolerated on 10G and BW25113, but not tolerated on BL21 (Figure 3A). Overall, these host-specific substitution patterns reveal a nuanced relationship between the tip domain composition and receptor preferences.

**Figure 3.**
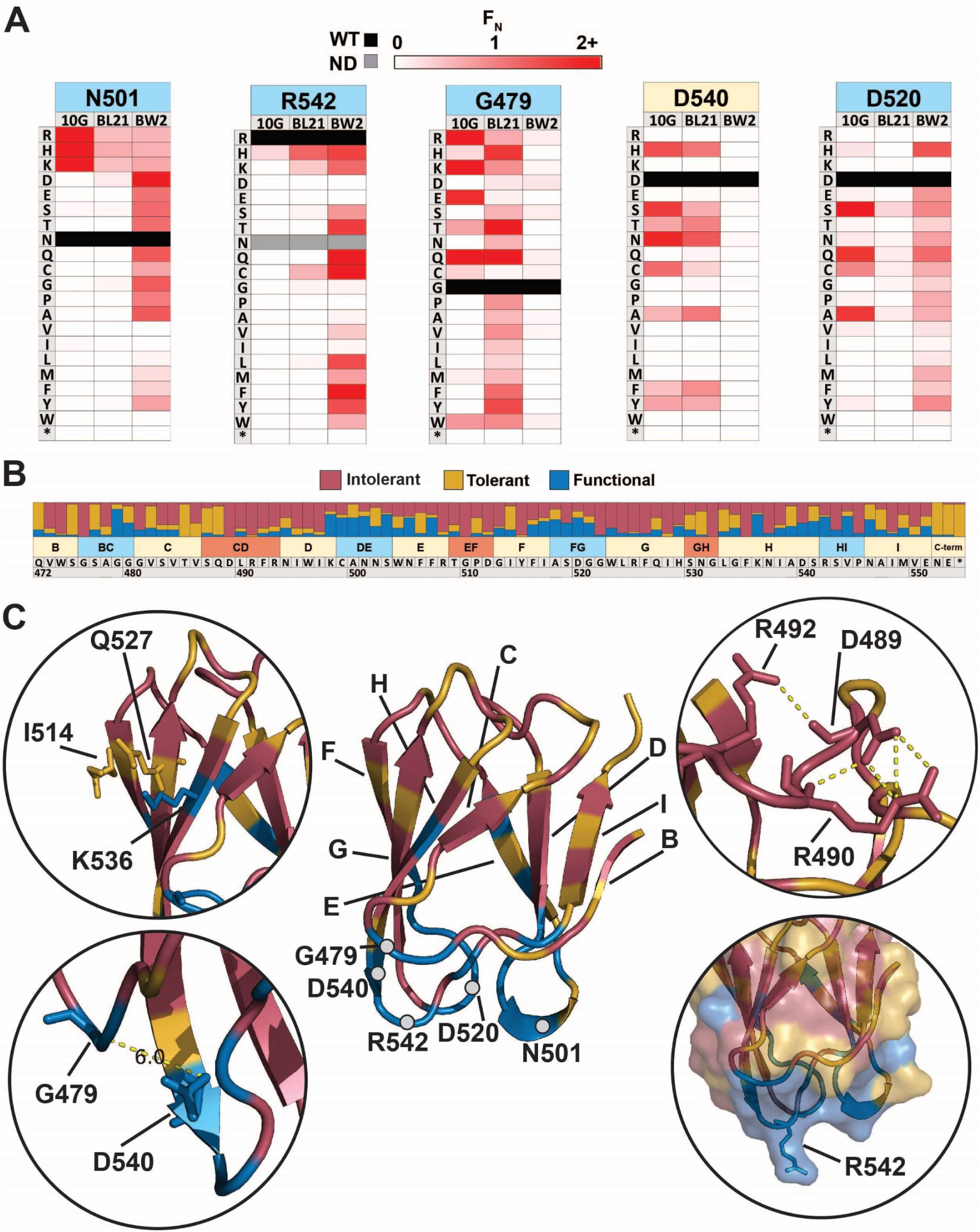
Comparison across hosts reveals regions of functional importance. **(A)** Host-specific differences in substitution patterns at five positions in the tip domain recapitulated from Figure 2. **(B)** Role of each position determined by aggregating scores of all substitutions in all hosts at that position. Substitutions are classified as intolerant (F_N_ < 0.1 in all hosts), tolerant (F_N_ ≥ 0.1 in all hosts), or functional (F_N_ < 0.1 in one host, F_N_ ≥ 0.1 in another host) **(C)** Crystal structure of the tip domain (center) with each residue colored as intolerant, tolerant, or functional based on the dominant effect at that position, β-sheets and residues listed in **(A)** are labeled. Key interactions defining function and orientation are highlighted in peripheral panels. See also Figure S5 and Table S4 and S5.

We quantitatively characterized the role of every residue by integrating selection data across all hosts to reveal a functional map of the tip domain at granular resolution (Figure 3B and 3C, Table S4). We classified every residue as ‘intolerant’, ‘tolerant’ or ‘functional’ based on aggregated F_N_ scores of all substitutions across all three hosts at every residue. Residues where the majority of substitutions were depleted were considered intolerant to substitution, while residues where at least a third of substitutions were depleted in one host and tolerated or enriched in another host were considered functional; the remaining positions were considered tolerant (see methods). The hydrophobic core comprising W474, I495, W496, I497, Y515, W523, L524, F526, I528, F535 and I548 is essential for stability and therefore is highly intolerant to substitutions (Figure 3B). Other intolerant positions include an elaborate network of salt bridge interactions involving D489, R491, R493, R508 and D512 in the interior loops, which likely constrain the orientation of the tip domain relative to the shaft (Figure 3C). Glycines generally provide conformational flexibility between secondary structure elements and normally tend to be mutable. Interestingly, several glycines (G476, G510, G522 and G532) are highly intolerant to substitutions. These glycines may be essential to minimize steric obstruction to adjacent larger residues, similar to G479 and D540 on BW25113 (Figure 3C). For example, G510 and G532 may facilitate formation of salt bridges in the interior loop, while G476 and G522 may facilitate a required receptor interaction in exterior loops for all three hosts.

It has been previously assumed that exterior loops are the primary functional region of the tip domain [25], [38]. We found that functional positions did typically point outward and are densely concentrated along exterior loops BC, DE, FG, and HI, as well as adjacent β-sheet residues. This is consistent with two specificity switching substitutions found in a previous study, D520Q and V544A, which are both located in exterior loops [41]. However, several residues in exterior loops, such as G476 and S543, were notably intolerant, indicating these residues may be poor targets for engineering or future combinatorial studies. Functional positions were also found in regions other than exterior loops, such as I514, Q527 and K536 which are β-sheet residues located along one side of the tip domain (Figure 3C). This suggests the phage can use the ‘side’ of the tip domain to engage the receptor, increasing the apparent functional area of the tip domain and highlighting several new regions as valuable engineering targets.

We also determined if the functionally important regions could be predicted computationally, as the ability to predict functionally important regions without DMS could rapidly accelerate engineering efforts. We used Rosetta, a state-of-the-art protein modeling software, to calculate the change in Gibbs free energy (ΔΔG) for each of the 1660 mutations and compared this distribution to our DMS results (Figure S5, Table S5, see methods). Predicted thermodynamic changes in stability mapped very well with over 93% of tolerated or functional positions predicted to have favorable ΔΔG energies. Incorporating stability estimations could further improve the engineering power of the assay. For example, substitutions predicted to be stable but that are intolerant in the DMS assay may indicate that the substituted residue is necessary for all three hosts.

Overall, these results paint a complex enrichment profile for each host with some broad trends but subtle host-specific effects. These results suggest that exterior loops and some outward facing positions in β-sheets act as a reservoir of function-switching and function-enhancing mutations, likely promoting host-specific and orientation-dependent interactions between phage and bacterial receptors. Functional positions identified by this comparison are ideal engineering targets to customize host range and activity.

### Discovery of gain-of-function variants against resistant hosts

The tail fiber is considered a reservoir of gain-of-function variants due to its principal role in determining fitness of a phage through host adsorption [22], [25]. We hypothesized that novel gain of function variants against a resistant host could be discovered by subjecting our tail fiber variant library to selection on a resistant host. To identify a resistant host, we focused on host genes *rfaG* and *rfaD* involved in the biosynthesis of surface lipopolysaccharide (LPS) which is a known receptor for T7 in *E. coli* [27]–[29]. Gene *rfaG* (synonyms *WaaG* or *pcsA*) transfers glucose to the outer core of LPS and deletion strains lack the outer core of LPS [43], while *rfaD* (synonyms *gmhD* or *WaaD*) encodes a critical epimerase required for building the inner core of LPS [44] (Figure 4A). Deletion of either gene reduces the ability of T7 to infect *E. coli* by several orders of magnitude (Figure 4F). We challenged the library of T7 variants against *E. coli* deletion strains BW25113Δ*rfaG* and BW25113Δ*rfaD* through pooled selection and deep sequencing as before (Figure 2), and determined a F_N_ score for each substitution on both strains (Figure 4B-C, Table S2 and S3). Independent replicates showed good correlation for BW25113Δ*rfaG* (R=0.99, 0.93, 0.93) but only adequate correlation for BW25113Δ*rfaD* (R=0.51, 0.68, 0.39) (Figure S3E-F). Although the scale of F_N_ was inconsistent across replicates on BW25113Δ*rfaD*, the same substitutions were largely enriched in all three replicates, suggesting reproducibility of results. Inconsistencies in F_N_ scores may arise due to severe loss of diversity causing stochastic differences in enrichment to become magnified across independent experiments.

**Figure 4.**
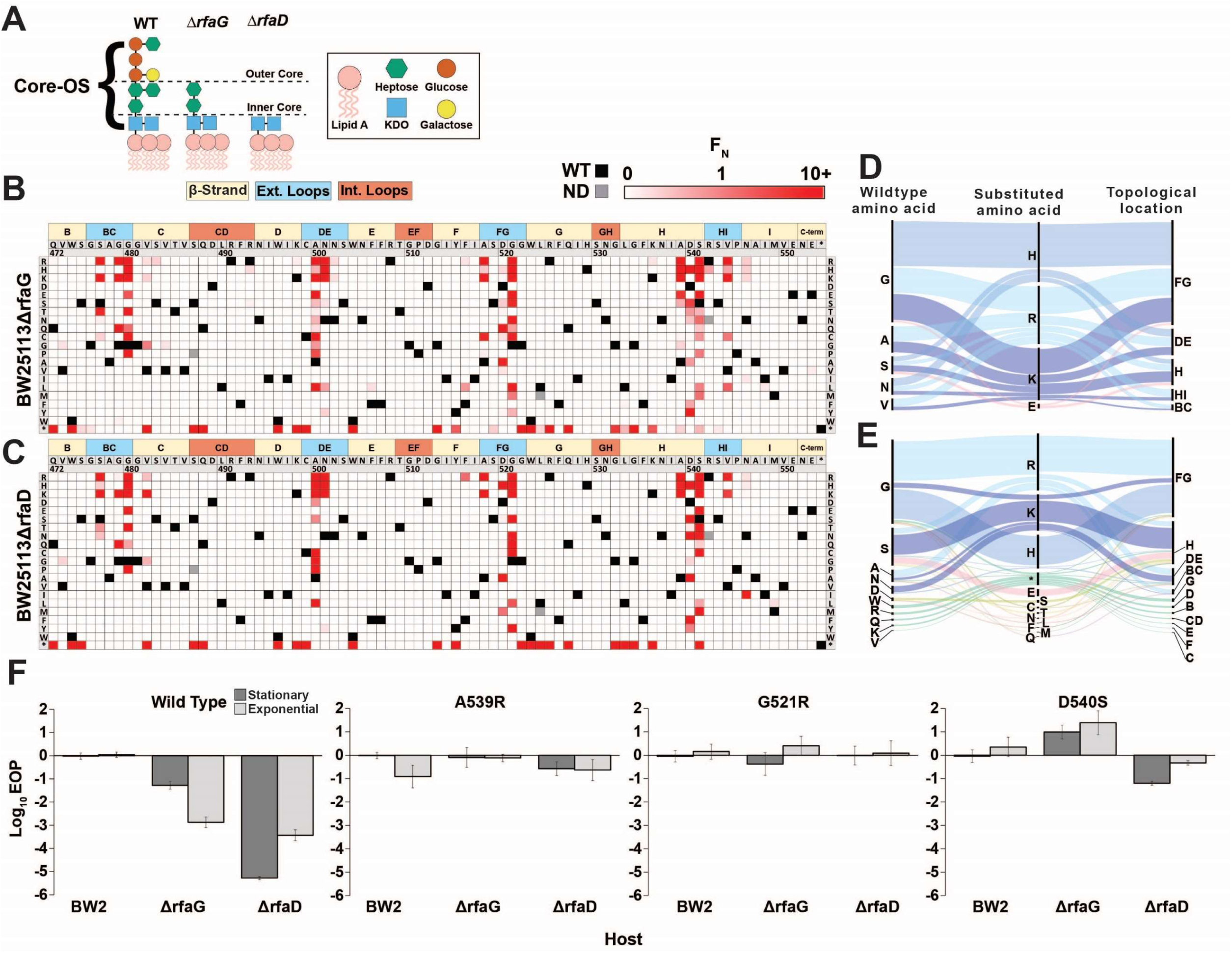
Discovery of gain-of-function variants against resistant hosts. **(A)** Schematic view of the LPS on wildtype BW25113, BW25113Δ*rfaG* and BW25113Δ*rfaD*. **(B-C)** Heat maps showing normalized functional scores (F_N_) of all substitutions (red gradient) and wildtype amino acid (solid black) at every position for BW25113Δ*rfaG* **(B)** and BW25113Δ*rfaD* **(C) (D-E)** Among highly enriched variants (F_N_ ≥ 10), targeted amino acids (left), their substitutions (middle) and topological location on the structure (right) on BW25113Δ*rfaG* **(D)** and BW25113Δ*rfaD* **(E)**, with each alluvial colored based on the substituted amino acid and scaled by F_N_. **(F)** EOP (mean ± SD, biological triplicates) for wildtype and select variants on BW25113 (BW2), BW25113Δ*rfaG* and BW25113Δ*rfaD* in exponential (dark gray) and stationary phases (light gray) using BW25113 as a reference host. See also Figure S3 and S6 and Table S3.

We engineered several gain-of-function T7 variants that could infect both deletion strains with activity comparable to wildtype T7 infecting susceptible BW25113 (Figure 4F). Low sequence diversity and high enrichment scores of T7 variants indicates a strong selection bottleneck which is consistent with diminished activity of wildtype T7 on the deletion strains. This is reflected in the significantly lower functional score of wildtype T7 on BW25113Δ*rfaG* and BW25113Δ*rfaD* (F=0.09 and F=0.03, respectively) in comparison to BW25113 (F=2.26) (Table S2). The number of enriched variants outperforming wildtype T7 (F_N_≥2) on the deletion strains (BW25113Δ*rfaG*: 55 variants, 3.3% and BW25113Δ*rfaD*: 68 variants, 4.1%) was over 3 times higher than BW25113 (16 variants, 1%) but comparable to 10G (48 variants, 2.9%) (Figure S6A-B). However, the enrichment scores of top performing variants such as G521H and G521R on BW25113Δ*rfaG* and S541K and N501H on BW25113Δ*rfaD* was over 100 times greater than wildtype T7, suggesting strong gain-of-function on the deletion strains (Figure S6C and Table S3). Of the 78 variants with F_N_≥2 on either deletion strain, 45 variants had F_N_≥2 on both strains indicating that variants that performed well on one strain typically performed well on the other strain (Table S3). This implies the enriched variants may have broad affinity for truncated LPS but cannot discriminate based on the length of the LPS. Nonetheless, hydrophilic substitutions were more strongly enriched on *BW25113ΔrfaG* (p<0.001), but not as significantly on *BW25113ΔrfaD* (p<0.033), suggesting subtle differences in surface chemical properties of deletion strains leading to host-specific enrichments (Figure S6D-F). Indeed, there were several variants with contrasting F scores on both strains such as S541T (*BW25113ΔrfaD* F_N_ = 44.8, *BW25113ΔrfaG* F_N_ = 0.6) and G521E (*BW25113ΔrfaD* F_N_ = 0, *BW25113ΔrfaG* F_N_ = 17.4) suggest potential host preference. Most substitutions were concentrated in the exterior loops BG, FG, HI, and β-strand H, all pointing downwards towards the bacterial surface, reinforcing the functional importance of these regions of the tip domain (Figure 4D and 4E). Notably, the most enriched variants had large positively charged substitutions (K, R, and H) akin to the enrichment pattern on 10G, suggesting the bacterial surface of these truncated mutants likely resembles that of 10G. Our results are consistent with a recent continuous evolution study, which identified G480E and G521R as possible gain-of-function variants on a strain similar to *BW25113ΔrfaD* and G479R and G521S as possible gain of function variants on *BW25113ΔrfaG* [22], although these variants only represent a small fraction of the gain of function variants discovered in our study.

We validated the results of the pooled selection experiment by clonally testing the ability of phage variants with high F_N_ (A539R, G521H, and D540S) to plaque on the deletion strains based on a standard efficiency of plating assay (EOP). Indeed, EOP results showed significant gain of function in these variants on the deletion strains (Figure 4F). D540S was particularly noteworthy as it performed better on the deletion strain *BW25113ΔrfaG* over wildtype BW25113 by 1-2 orders of magnitude. Based on these results, we conclude that D540 is critical for infecting wildtype BW25113 (Figure 3) likely by interacting with the outer core of LPS. When the outer core of the LPS is missing (*BW25113ΔrfaG*), a substitution at this position becomes necessary for adsorption either to a different LPS moiety or to an alternative receptor.

We introduced stop codon at every position to systematically evaluate the function of tip domains truncated to different lengths. Many truncated variants performed well, especially on BW25113Δ*rfaG* which included some with F_N_≥10 (Table S3). We clonally tested variant R525*, the best performing truncated library member (BW25113Δ*rfaG* F_N_ = 9.55, BW25113Δ*rfaD* F_N_ = 75.7), and found that this mutant showed no ability to plaque on any host unless provided the tail fiber *in trans*. These truncated phages, detectable here only using deep sequencing, may demonstrate how obligate lytic phages could become less active in a bacterial population, slowly replicating alongside their bacterial hosts, requiring only a single mutation to become fully active again. In fact, acceptor phages altogether lacking a tail fiber were present at extremely low abundance (Table S2). These phages are not artifacts from library creation as some ability to replicate is required to produce detectable concentrations of each phage. We concluded that these are viable phage variants albeit with a much slower infection cycle resulting in their inability to form visible plaques.

### Targeting pathogenic *E. coli* causing urinary tract infection using T7 variants

Phage therapy is emerging as a promising solution to the antibiotic resistance crisis. Recent clinical success stories against multidrug resistant *Acinetobacter* and *Mycobacterium* showcase the enormous potential of phage therapy [4], [9]. However, development of effective phage-based therapeutics is hindered by low phage susceptibility, resulting in rapid onset of bacterial resistance. Although initial application of phages in a laboratory setting may reduce bacterial levels, the residual bacterial load remains high, causing bacteria to quickly recover after phage application [45]–[47]. A high ratio of phage to bacteria (multiplicity of infection or MOI) may productively kill bacteria in a laboratory setting by overwhelming a host with many phages [48]. However, ensuring an overwhelming amount of phages in a clinical setting is not always feasible [49]. Engineering highly active phages that overcome bacterial resistance and can therefore productively eliminate bacterial populations at low MOI in a laboratory setting would greatly enhance phage-based therapeutics. We hypothesized that engineered tip domain variants may abate bacterial resistance and be active even at low MOI by better adsorbing to the native receptor or recognizing a new bacterial receptor altogether.

To test this hypothesis, we chose a pathogenic *E. coli* strain isolated from a patient with urinary tract infection (UTI) [50]. Although T7 can infect the UTI strain, resistance arises rapidly, a phenomenon all too common with the use of natural phages. EOP assays for wildtype T7 showed resistant plaque morphology and no visible lysis was detected when wildtype T7 was applied in liquid culture (MOI=1), indicative of rapidly emerging resistance. However, the variant library applied on the UTI strain cleared the culture (MOI=1) suggesting T7 variants are capable of lysing and attenuating resistance exist in the pool. We clonally characterized three variants (N501H, D520A and G521R) isolated from plaques. All three variants vastly outperformed wildtype T7 in terms of onset of resistance. Resistance emerged approximately 11-13 hours after initial lysis for the three variants whereas it took merely 1-2 hours for wildtype (Figure 5A). In particular, the N501H variant lysed cells faster and produced a lower bacterial load post lysis, suggesting far greater activity compared to wildtype T7. Next, we compared the effect of phage MOI (MOI=10^2^-10^-5^) on the lysing activity of N501H and wildtype T7 (Figure 5B). At lower MOI, time to lysis of N501H was half that of wildtype T7, though they were comparable at higher MOI.

**Figure 5.**
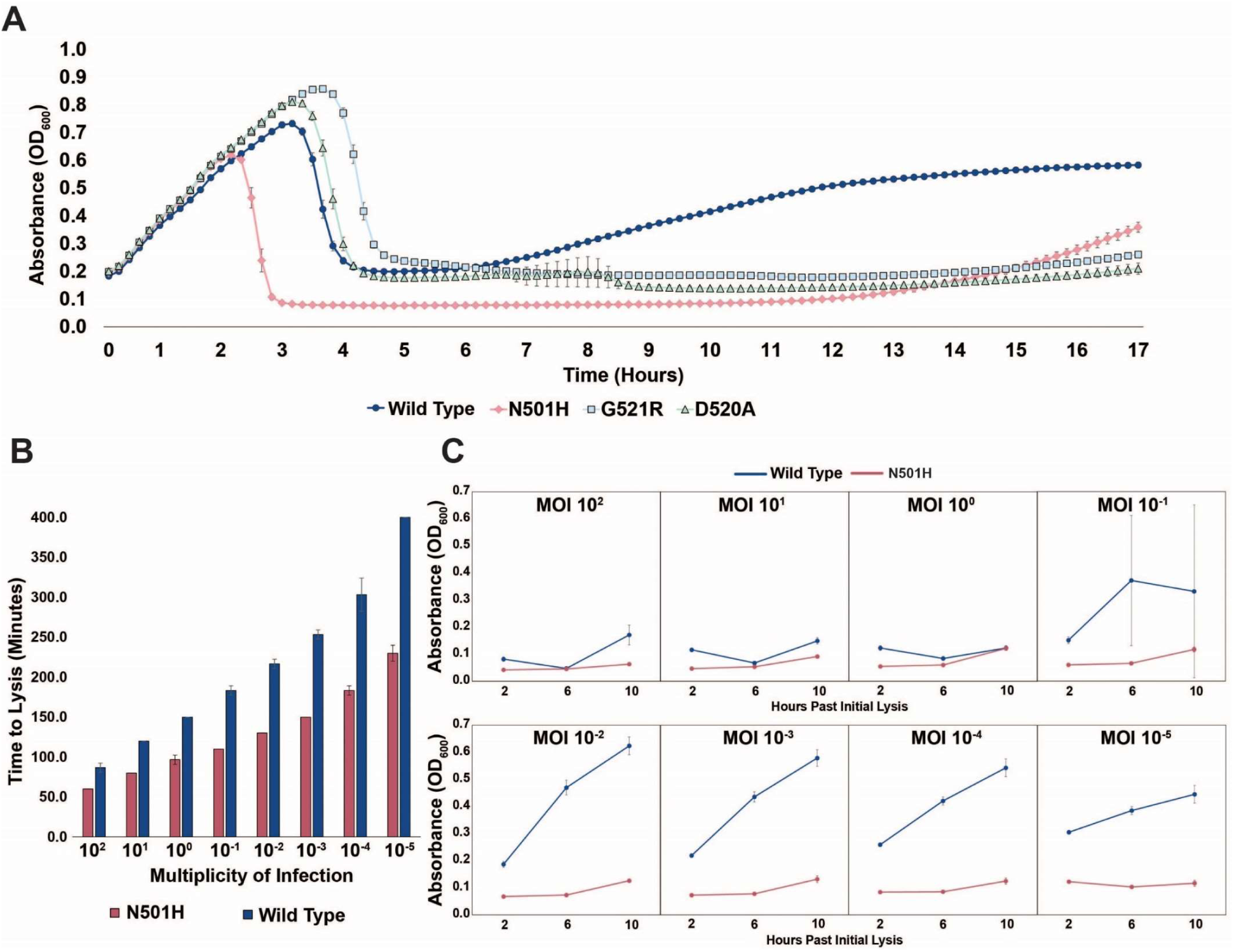
Targeting pathogenic *E. coli* causing urinary tract infection using T7 variants. **(A)** Growth time course of UTI strain subject to wildtype T7 and select variants. Phages were applied after an hour at an MOI of ∼10^-2^ **(B)** Estimated time to lysis of UTI strain incubated with wildtype T7 and N501H variant over a range of MOIs, derived from time course experiments. **(C)** Cell density (OD600) of UTI strain when incubated with wildtype T7 and N501H variant at select timepoints after initial lysis. All data represented as mean ± SD of biological triplicate.

A striking contrast between N501H and wildtype T7 is evident in the suppression of resistance at progressively lower MOI (Figure 5C). Between MOI of 100 to 1, both N501H and wildtype perform similarly. However, between MOI of 10^-1^ and 10^-5^, N501H suppresses resistance over a 10-hour window, while wildtype phages are quickly overcome by the bacterial resistance. We postulate that at high MOI wildtype T7 simply overwhelms the host before resistance arises due to multiple infection events happening simultaneously, a known outcome for high MOI conditions [48]. At lower MOI, resistance can emerge, and only phages adapted to the host can effectively kill the host. These results indicate that ORACLE can generate phage variants superior to wildtype phage that could then become starting points for further engineering therapeutic phages.

### Host range constriction emerges from global comparison across variants

Most phages are specialists that selectively target a narrow range of hosts but avoid other closely related hosts [51]. We wanted to assess differences in the host range of individual variants on 10G, BL21 and BW25113 and identify variants with constricted host ranges. Ideally, host specificities can be determined by subjecting a co-culture of all three hosts to the phage library. However, deconvolving specificities of thousands of variants from a pooled co-culture experiment can be technically challenging. Instead, we sought to estimate specificities by comparing F_N_ of a phage variant on all three hosts.

Although F_N_ compares activity of variants within a host, it could nonetheless be a useful proxy for estimating specificities across hosts. For instance, a phage variant with high F_N_ on BL21 but completely depleted on BW25113 is more likely to specifically lyse BL21 than BW25113 in a co-culture experiment. Based on this rationale, we considered different metrics of comparison of F_N_, and settled on difference in F_N_ of a variant with reduced weight for enrichment (or F_D,_ see methods) between any two hosts as an approximate measure of host preference. This metric is not an absolute measure of host specificity, but one devised to reveal broad trends in specificity to prioritize variants for downstream validation.

To assess if variants preferred one host over another, we computed F_D_ for all three pairwise combinations and plotted functional substitutions as points on or above/below a ‘neutral’ line (Figure 6A-C, Table S4). Variants above the line favor lysis of the noted host, and vice versa for variants below the line. To check if this F_D_-based approach is suitable for assessing host specificity, we compared our results with previously published data. Two substitutions, D520Q and V544A, that were reported to have a preference for BW25113 and BL21, respectively, in head to head comparisons [41] were placed correctly in our plots, confirming the validity of our F_D_-based classification scheme. We identified 118 out of 1660 variants as good candidates for constricting host range (|F_D_|≥1). Of the 118 variants, 53 variants favor BW25113 over BL21 and 98 variants favor BW25113 over 10G in pairwise comparisons (Figure 6A and 6C, Table S4). Between BL21 and 10G, there are 15 variants that favor BL21 but none that favor 10G (Figure 6B, Table S4).

**Figure 6.**
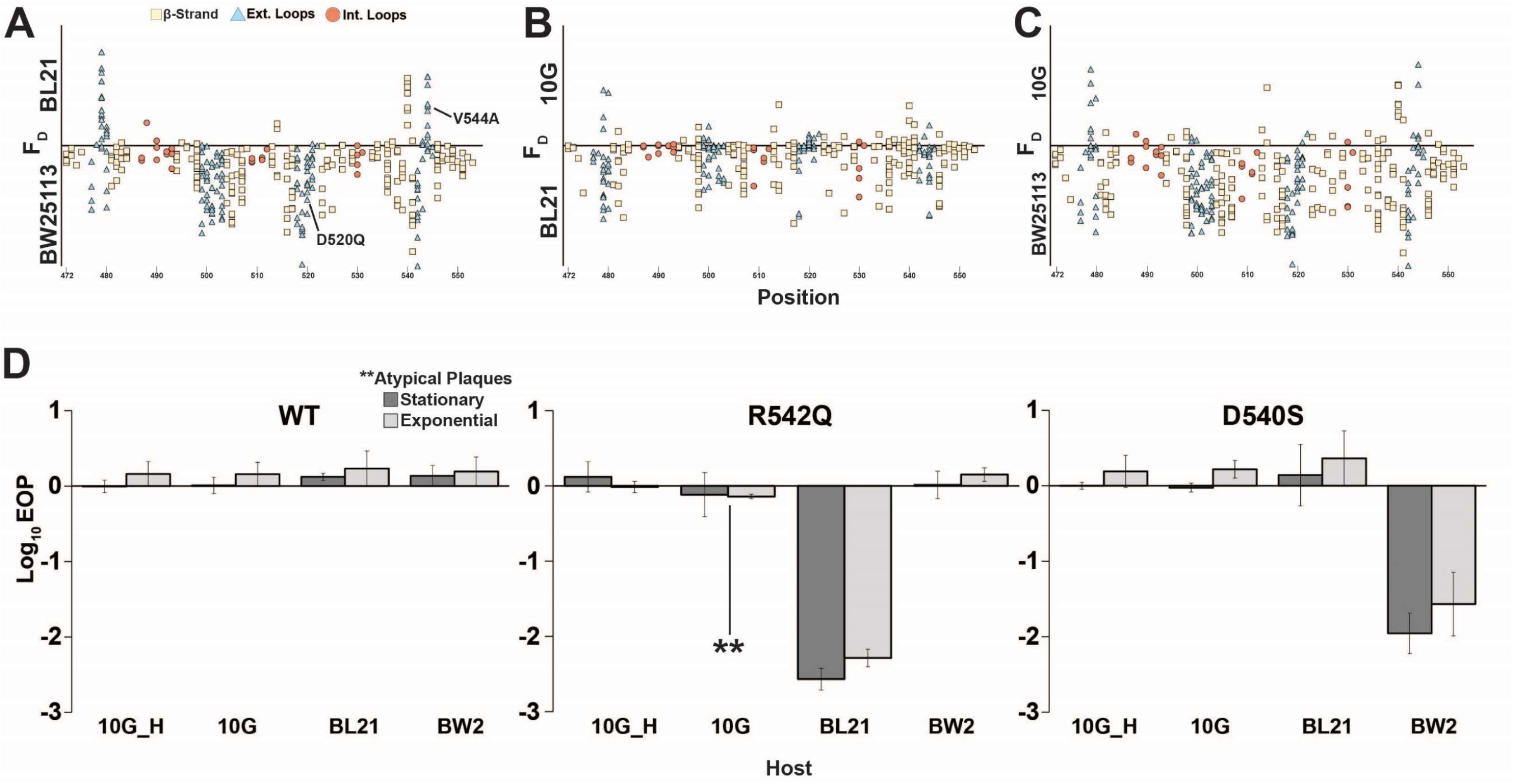
Host range constriction emerges from global comparison across variants. **(A-C)** Pairwise comparison of differences in functional scores of variants between hosts (see Methods). Variants above the line favor lysis of host noted above the line, and vice versa for variants below the line. **(D)** EOP (mean ± SD, biological triplicates) for wildtype T7 and select variants on BW25113, 10G and BL21 in exponential (dark gray) and stationary phases (light gray) using exponential 10G with *gp17* tail fiber helper plasmid (10G H) as a reference host. R542Q plaques are atypically small until EOP ∼10^-2^. See also Table S5.

Certain key positions including G479, D540, R542 and D520 which we previously identified as functionally important (Figure 3A) are the molecular drivers of specificity between hosts (Figure. 6A-C). Taken together, our data suggests that it would be easier to find a variant capable of specifically lysing BW25113, less so for BL21, and most challenging for 10G.

To validate our analysis, we clonally tested variant R542Q which had a greater preference for BW25113 than BL21 or 10G (BW25113 F_D_ =2.0, BL21 F_D_ =0, 10G F_D_=0) and variant D540S which had a greater preference for BL21 and 10G than BW25113 (10G F_D_ = 1.03, BL21 F_D_= 0.62, BW25113 F_D_ = 0.03). Indeed, R542Q showed a ∼100-fold decrease in the ability to plaque on BL21 compared to BW25113 while 10G plaques were atypically small, indicating a severe growth defect (Figure 6D). In contrast, D540S showed a ∼100-fold decrease in the ability to plaque on BW25113 compared to BL21 and 10G (Figure 6D), confirming the host constriction properties of these variants. In summary, pairwise comparison is a powerful tool to map substitutions that constrict host range and can be leveraged to tailor engineered phages for targeted hosts.

## Discussion

In this study, we used ORACLE to create a large, unbiased library of T7 phage variants to comprehensively characterize the mutational landscape of the tip domain of the tail fiber. Our study identified hundreds of novel function-enhancing substitutions that had not been previously characterized. We mapped regions of function-enhancing substitutions on to the crystal structure to rationalize how sequence and structure influence activity and host range. Several important insights emerged from these results. Cross-comparison between different hosts and selection on resistant hosts allowed us to map key substitutions leading to host discrimination and gain of function. Single amino acid substitutions are sufficient to enhance activity and host range including some that confer dramatic increases in activity or specificity. The functional landscape on each host is unique, reflecting both different molecular preferences of adsorption and the fitness of wildtype T7 on these hosts. For instance, hydrophilic substitutions were enriched in 10G while hydrophobic substitutions were enriched in BW25113. Notably, substitutions on 10G (an *E. coli* K-12 derivative lacking LPS components) mirrored substitutions that recovered function on BW25113 mutants with truncated LPS which shows convergence of selection. Function-enhancing substitutions were densely concentrated in the exterior loops indicating an orientational preference for receptor recognition. However, they were also found on other surface residues, albeit less frequently, suggesting alternative binding modes of the tip domain for host recognition, and several intolerant residues were located in exterior loops. Taken together, these results highlight the extraordinary functional potential of the tip domain and rationalizes the pervasive use of this structural fold in nature for molecular recognition. Comparison of these functional profiles precisely reveals regions that are ideal engineering targets for customizing host range and activity and identifies intolerant residues that should be avoided when engineering synthetic phages.

These results also highlight the power of deep sequencing to detect and resolve small functional effects over traditional low-throughput plaque assays. This is best illustrated in the case of truncated variants visible only to deep sequencing, but incapable of plaque formation without a helper plasmid. The truncated variants are likely not experimental artifacts as some ability to replicate is required to survive multiple rounds of selection on the host. Truncation of the tip domain may misorient the phage relative to the receptor, likely resulting in slower growth and deficiency in plaquing, while still capable of replicating. Since plaque formation is a complex process, inability to plaque may not imply a functionally incompetent phage.

ORACLE is designed as a foundational technology to elucidate sequence-function relationships in any phage gene. On T7, ORACLE can be used to investigate the function of several important genes including the remainder of the tail fiber and tail structure, capsid components, or lysins and holins. Together, these will provide a comprehensive view of the molecular determinants of the structure, function, and evolution of a phage. Once the phage variants are created, scaling up ORACLE to investigate potentially tens of hosts merely scales up sequencing volume, not experimental complexity. Such a large-scale study will lead to a detailed molecular understanding and adaptability of phage bacterial interactions. Any phage with a sequenced genome and a plasmid-transformable propagation host should be amenable to ORACLE because the phage variants are created during the natural infection cycle. This approach can be leveraged to tune activity for known phages with high activity, such as T7, or to identify engineering targets that dramatically increase activity for newly isolated natural phages.

The confluence of genome engineering, high-throughput DNA synthesis, and sequencing enabled by ORACLE together with viral metagenomics could transform phage biology. Phages constitute unparalleled biological variation found in nature and are aptly called the “dark matter” of the biosphere. Their sequence diversity and richness are coming to light in the growing volume of viral metagenome databases. However, what functions these sequences encode remains largely unknown. For instance, fecal viromes estimate 10^8^-10^9^ virus-like particles per gram of feces, but less than a quarter of sequence reads align to existing databases [52]. While this knowledge gap is daunting, it also presents an opportunity to mine metagenomic sequences to characterize their function and engineer programmable phages. By enabling sequence programmability, we envision ORACLE as a powerful tool to discover new phage ‘parts’ from metagenomic sequences.

## Author Contributions

P.H and S.R designed the study, analyzed the data, and wrote the manuscript.

P.H performed all experiments. A.M and P.H wrote the software. M.L and K.N. assisted with data analysis.

## Acknowledgements

We would like to thank Dr. Rodney Welch for UTI473 strain and Dr. Douglas Weibel for BW25113 deletion mutant strains. We thank Dr. Karthik Anantharaman, Dr. John Yin, Laura Alexander and Chutikarn Chitboonthavisuk for critical review of the manuscript. This work is partially supported by the US Department of Agriculture Hatch award (WIS02066) and the Gates Grand Challenges grant (OPP1150209). A.M is supported by the Great Lakes Bioenergy Research Center (U. S. Department of Energy Award Number DE-SC0018409). M.L is supported by NIH Molecular Biophysics Training Program T32 GM08293 and William H. Peterson Fellowship in Biochemistry. K.N is supported by NIH National Research Service Award T32 GM07215 and the Robert and Katherine Burris Biochemistry Fund.

---

**Figure S1.**
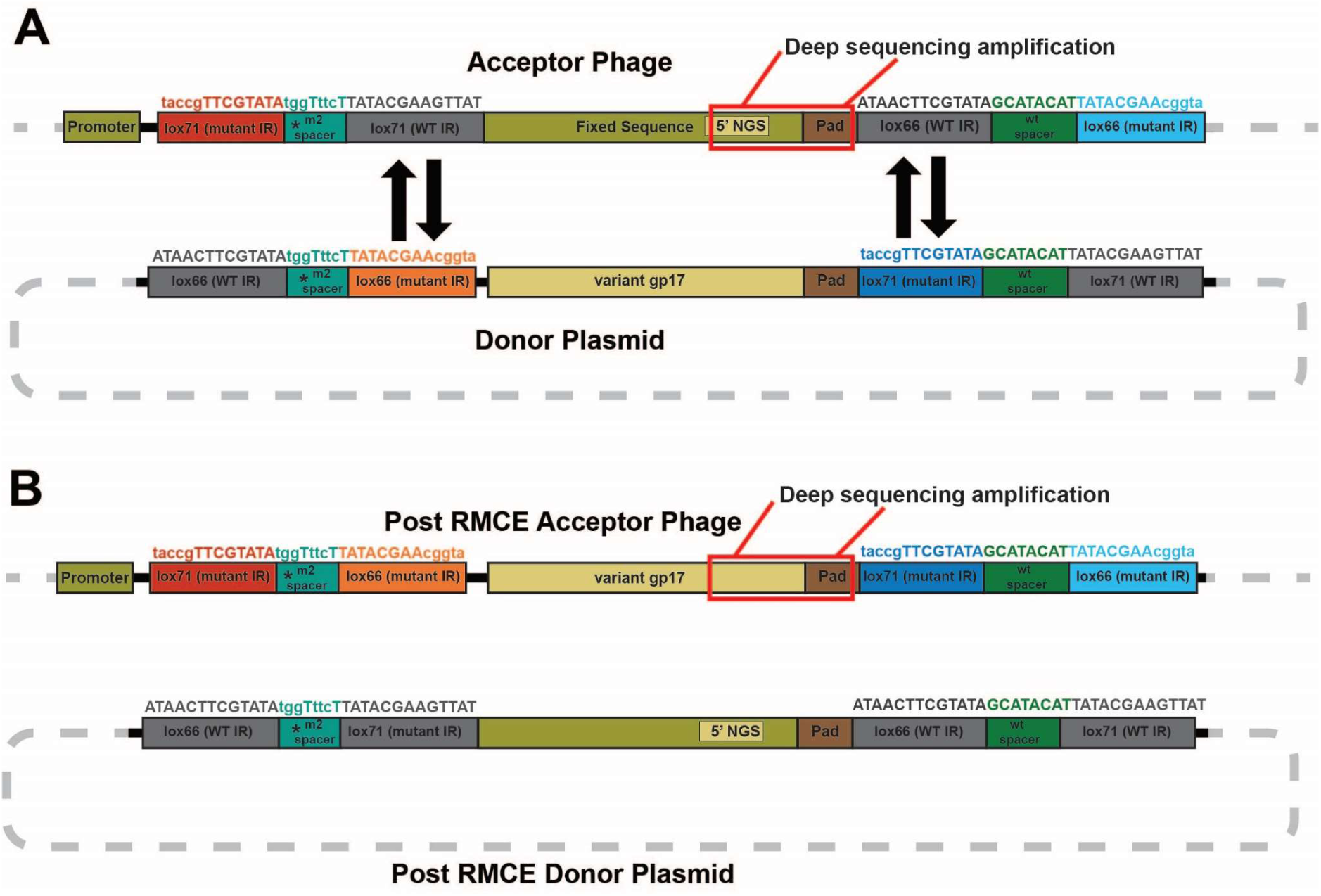
Sequence rearrangements before and after recombinase-mediated cassette exchange. Related to Figure 1. Schematic illustration of sequence rearrangements in acceptor phage and donor plasmid **(A)** before and **(B)** after recombinase-mediated cassette exchange (RMCE). Specific lox recombinase sites required for exchanging sequence cassettes (variant and fixed sequence) are shown. Lox sites have wild type (WT) or mutated inverted repeats (IR) and one-way RMCE can only occur if one IR is wild type, while the m2 spacer forces recombination in the correct orientation and prevents adverse recombination events [53]. Deep sequencing targets the area boxed in red between the 5’ NGS region and 3’ pad on both acceptor phages and the variant library.

**Figure S2.**
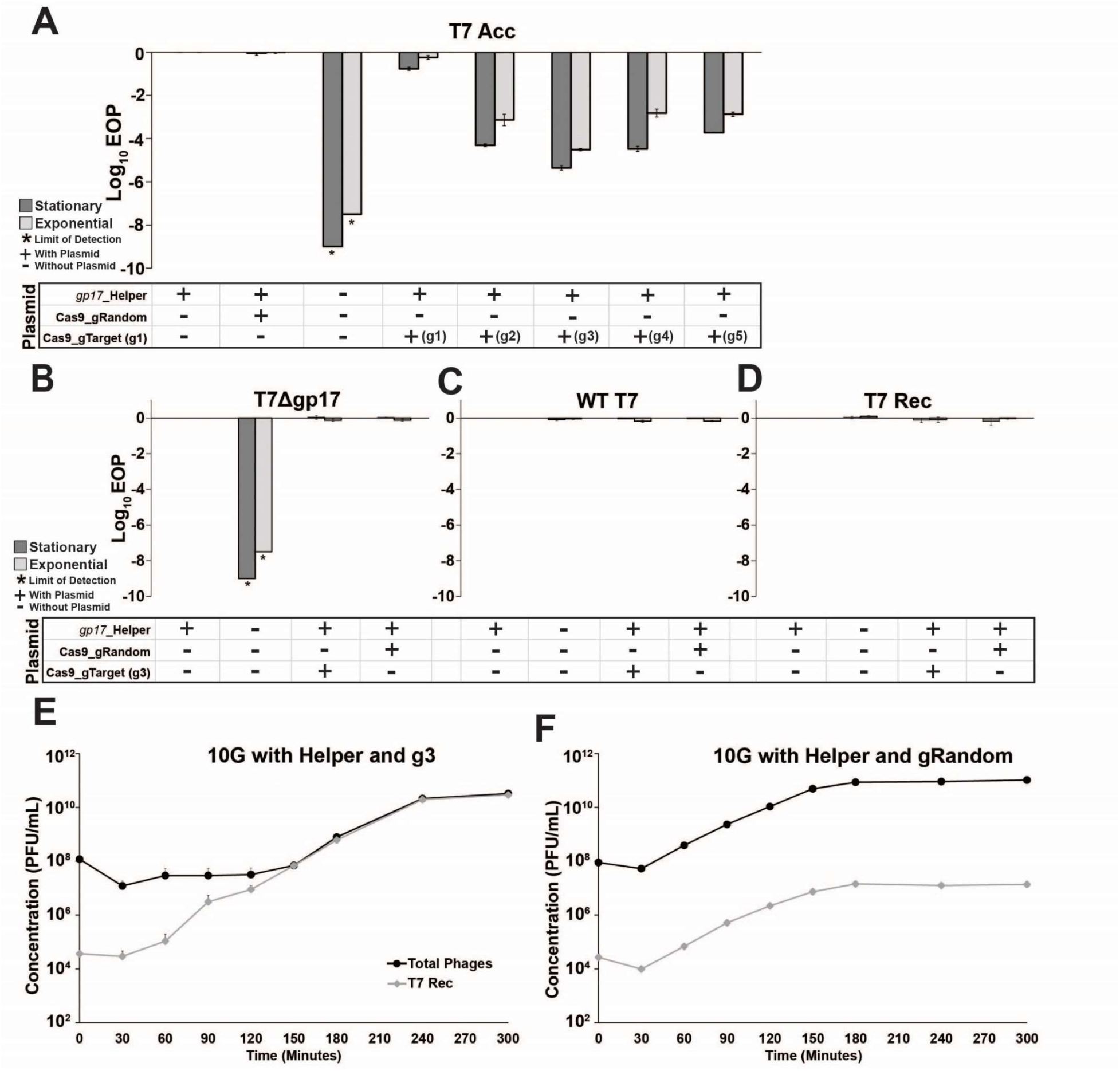
Effect of Cas9-gRNA system on acceptor and control phages. Related to Figure 1. **(A)** Efficiency of plating (EOP) measurements on *E. coli* 10G for T7 acceptor phage (T7 Acc) with combinations of *gp17* helper plasmid, Cas9-gRNA plasmid with a random guide (gRandom) or targeting guides 1 through 5 (gTarget, g1-g5). T7 acceptor phages cannot plaque without helper plasmid. Their plaquing is unaffected by Cas9-gRNA with a random guide(gRandom). Among five different gRNAs targeting fixed sequence, g3 shows highest targeting efficiency and was used for phage library construction with ORACLE. (**B-D)** The Cas9-gRNA system does not adversely affect the plaquing activity of untargeted phages. Efficiency of plating (EOP) measurements on *E. coli* 10G with combinations of *gp17* helper plasmid, Cas9-gRNA plasmid with a random guide or targeting guide 3 (gTarget, g3) for **(B)** T7 phage without *gp17* (T7*Δgp17*), **(C)** wildtype T7 (WT T7) and **(D)** acceptor T7 phage recombined with wildtype *gp17* (T7 Rec). **(E and F)** Comparison of accumulation of recombined phages (T7 Rec) with respect to total phages using **(E)** 10G with *gp17* helper plasmid and Cas9-gRNA (g3) and **(F)** 10G with *gp17* helper plasmid and Cas9-gRNA (gRandom). All data shown is biological triplicates (mean + SD), all EOP data uses 10G with *gp17* helper as a reference host.

**Figure S3.**
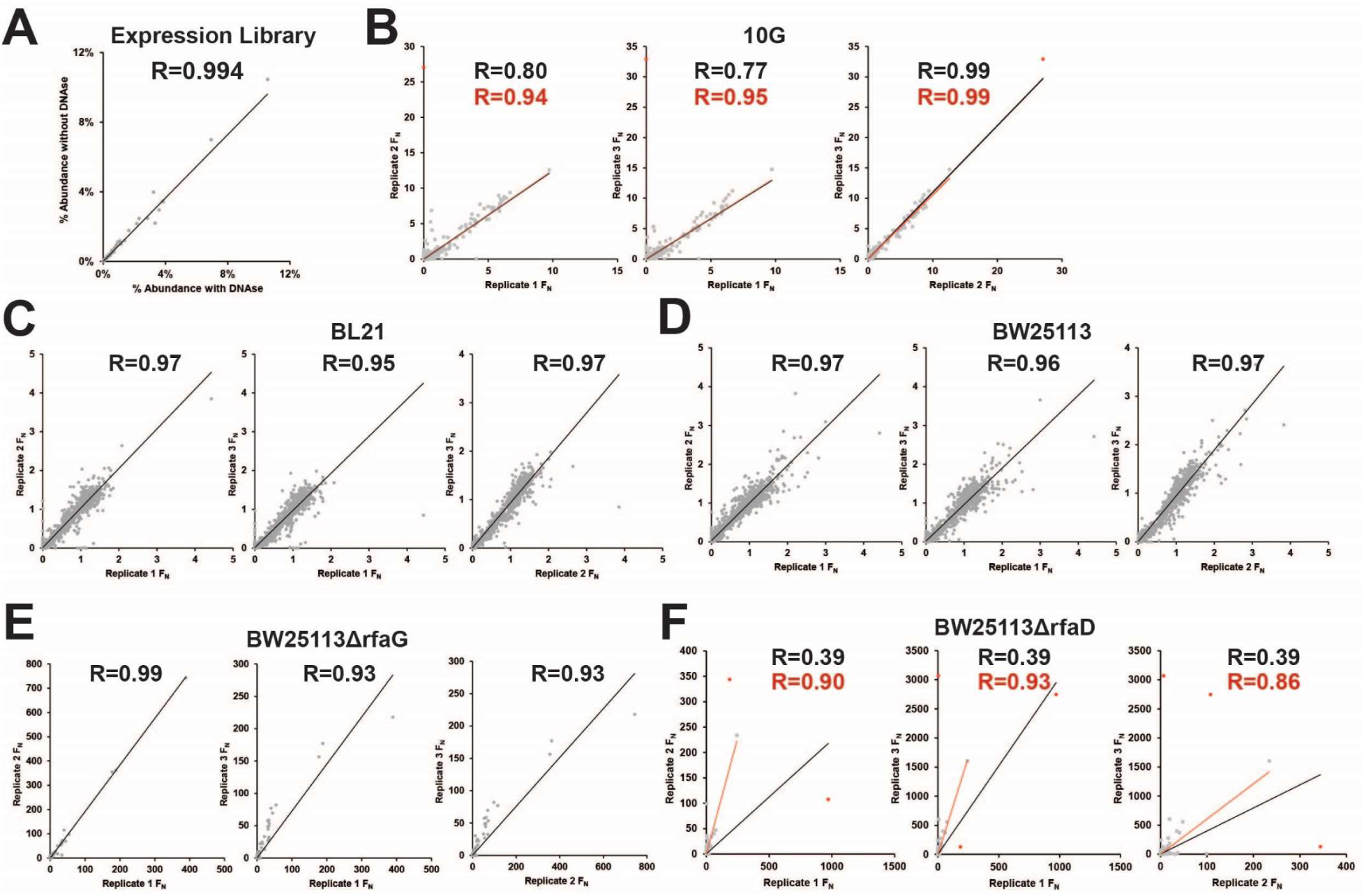
Correlation between biological replicates for selection of phage variant library under different conditions. Related to Figure 2 and 4. Correlation of F_N_ scores between biological replicates of phage variant library **(A)** with and without DNAse treatment and on multiple hosts and deletion mutants including (B) *E. coli* 10G **(C)** BL21 **(D)** BW25113 **(E)** BW25113Δ*rfaG* and **(F)** BW25113Δ*rfaD*. R values and trendlines are displayed for all variants (black) and with outliers excluded for 10G and BW25113Δ*rfaD* (red) with outliers in red and all other points in grey.

**Figure S4.**
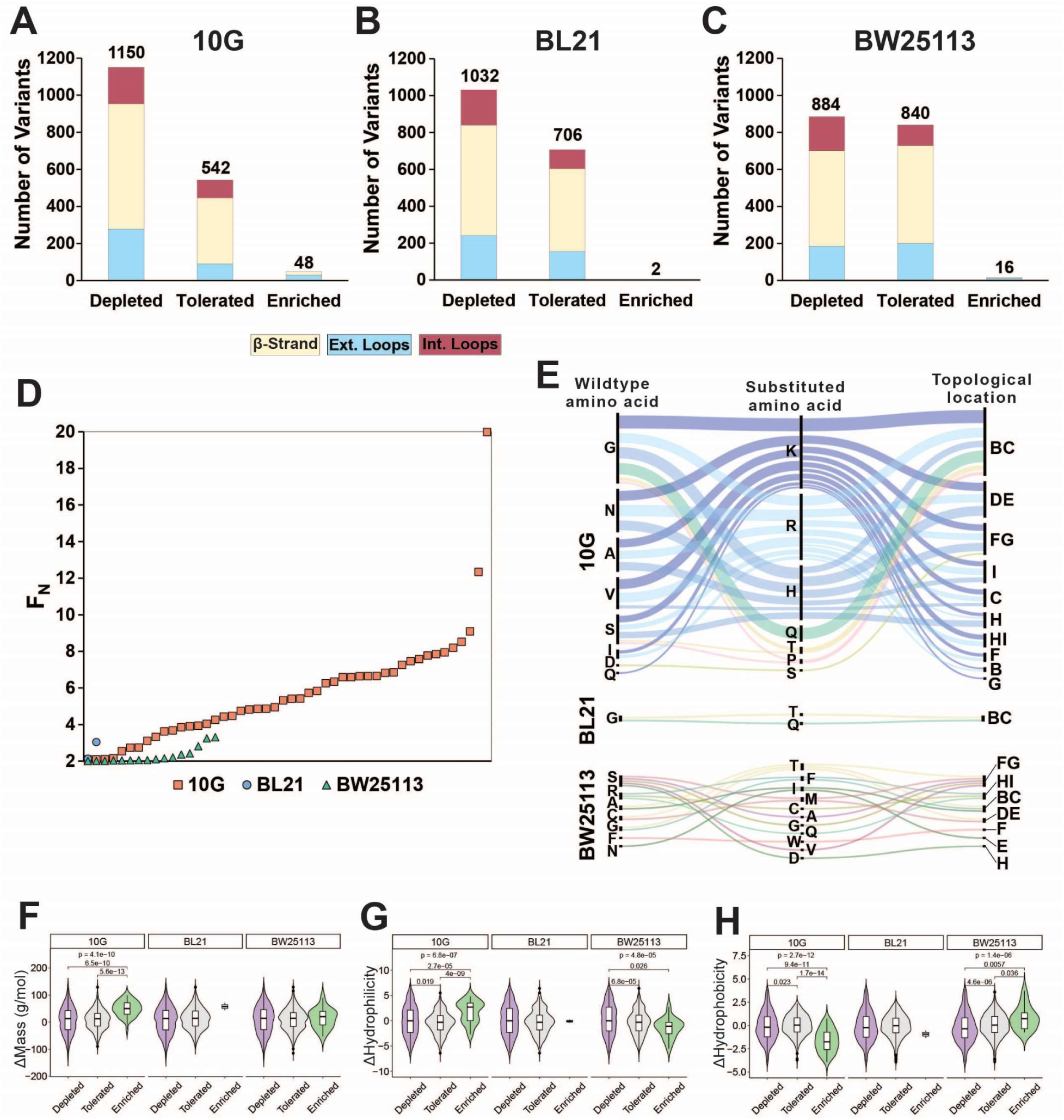
Distribution, enrichment profile and physicochemical properties of variants after selection on 10G, BL21 and BW25113. Related to Figure 3. Number of variants that were depleted (F_N_ ≤ 0.1) tolerated (F_N_ >0.1 and <2) or enriched (F_N_ ≥ 2) after selection on (A) *E. coli* 10G, **(B)**, BL21, and **(C)** BW25113, separated by topology of the tip domain color coded as interior loops (red), β-sheets (beige) and exterior loops (blue). **(D)** Average F_N_ of enriched variants (F_N_ ≥ 2) for 10G (orange squares), BL21 (blue circles), and BW25113 (teal triangles) ordered left to right from lowest to highest F_N_. **(E)** Alluvial distribution of enriched variants (F_N_ ≥ 2) on 10G (upper), BL21 (middle) and BW25113 (bottom), showing wild type amino acids (left), their substitution (middle) and topological location on the structure (right). Each alluvial is colored based on the substituted amino acid and scaled by F_N_ across hosts. Violin plots comparing **(F)** change in mass, **(G)** change in hydrophilicity, and **(H)** change in hydrophobicity for grouped depleted (F_N_ ≤ 0.1) tolerated (F_N_ >0.1 and <2) or enriched (F_N_ ≥ 2) substitutions on *E. coli* 10G, BL21 and BW25113. p-values are shown if only if <0.05, the upper p-value is the result of a Kruskal-Wallis test among all three groups while pairwise p-values from a Wilcoxon test are shown linking each group.

**Figure S5.**
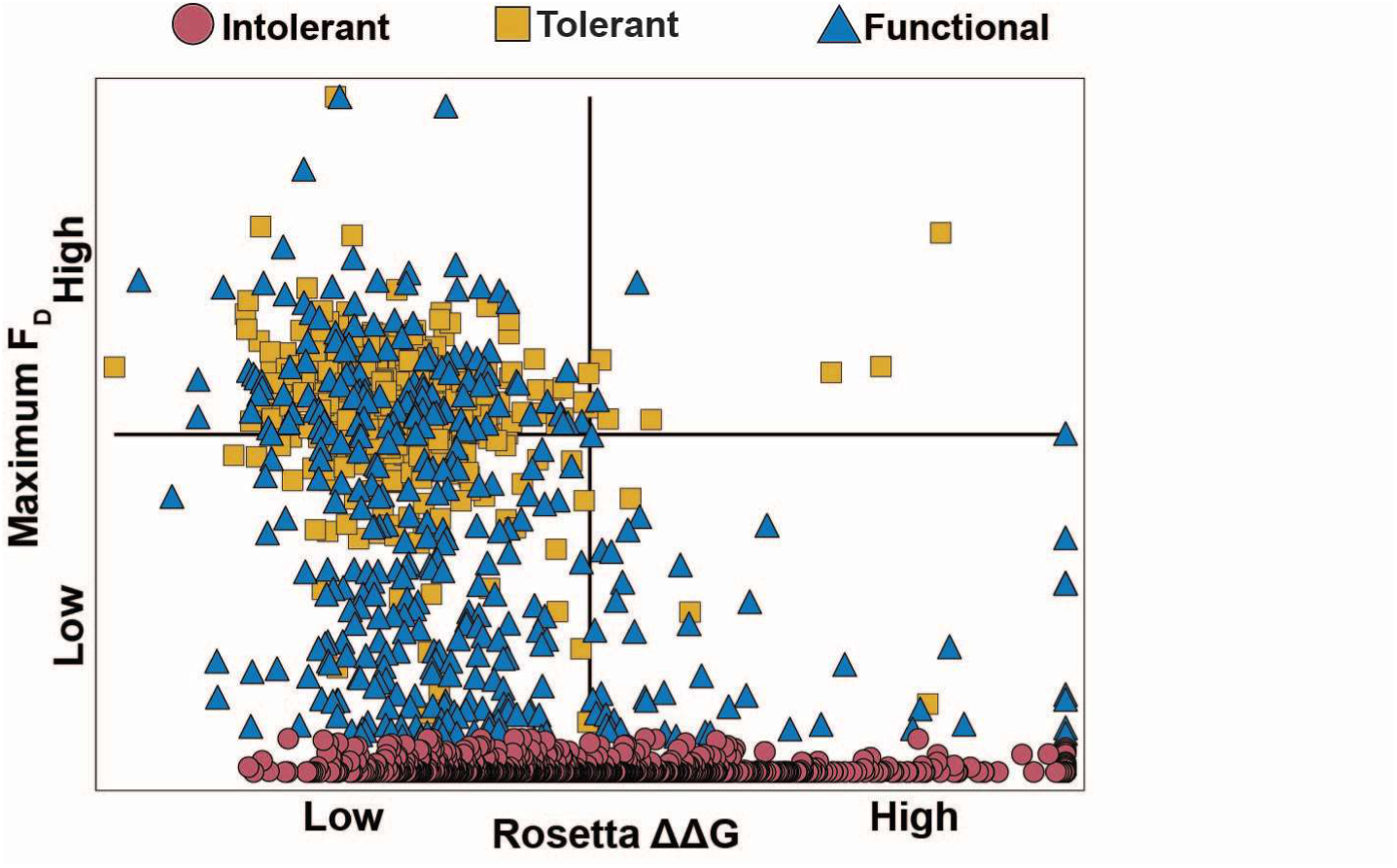
Comparing F_D_ to computationally predicted stability of all variants. ΔΔG. Related to Figure 3. Comparison of maximum F_D_ values between *E. coli* 10G, BL21 and BW25113 to computationally predicted change in stability (ΔΔG, see methods) for each intolerant (red circles), tolerant (yellow squares) and functional (blue triangles) in the variant library.

**Figure S6.**
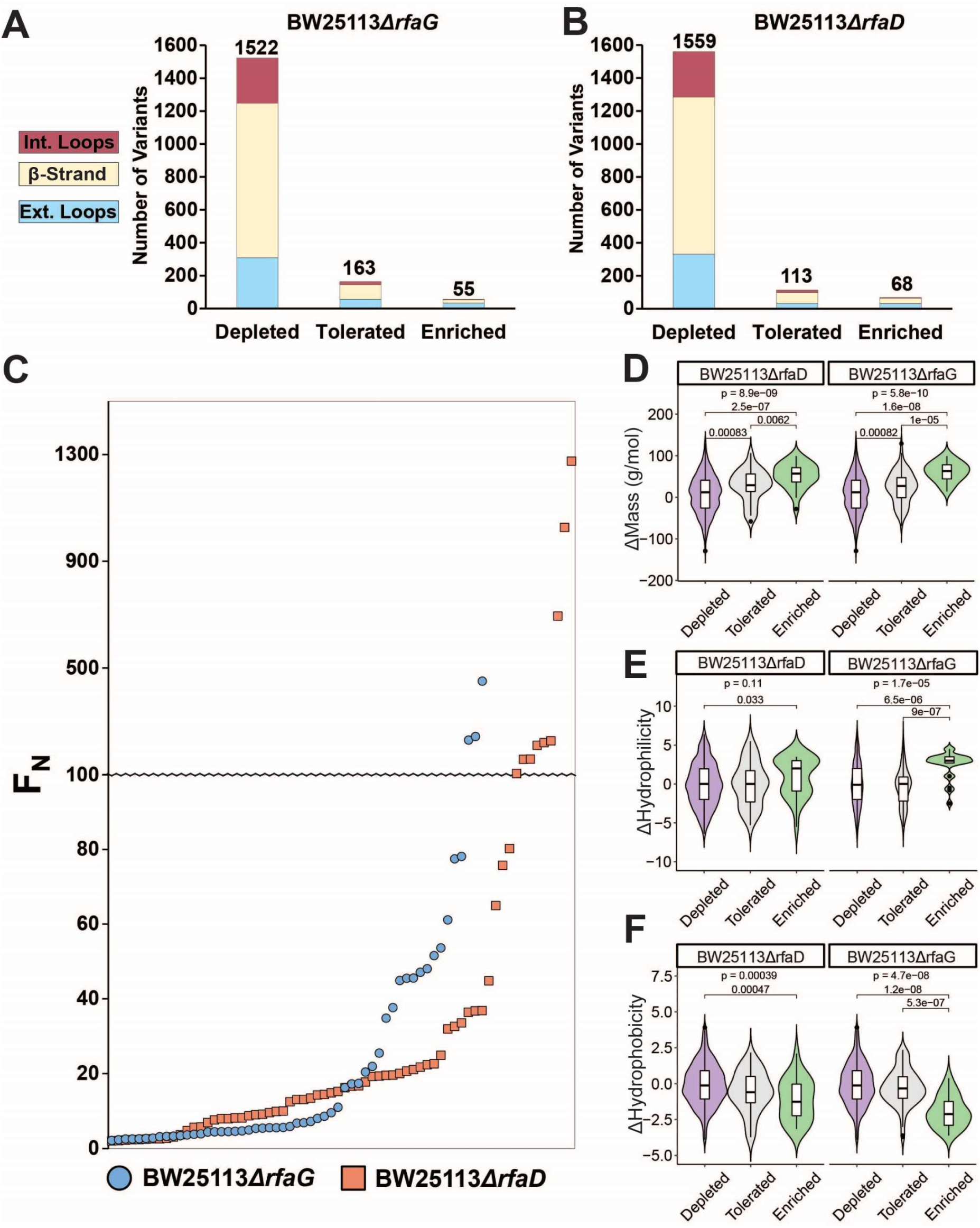
Distribution, enrichment profile and physicochemical properties of variants after selection on BW25113ΔrfaG and BW25113ΔrfaD. Related to Figure 4. Number of variants that were depleted (F_N_ ≤ 0.1) tolerated (F_N_ >0.1 and <2) or enriched (F_N_ ≥ 2) after selection on **(A)** BW25113*ΔrfaG* or **(B)** BW25113*ΔrfaD*, separated by topology of the tip domain color coded as interior loops (red), β-sheets (beige) and exterior loops (blue). **(C)** Average F_N_ of enriched variants (F_N_ ≥ 2) for BW25113*ΔrfaD* (orange squares) and BW25113*ΔrfaG* (blue circles) ordered left to right from lowest to highest F_N_. Violin plots comparing **(D)** change in mass, **(E)** change in hydrophilicity, and **(F)** change in hydrophobicity for grouped depleted (F_N_ ≤ 0.1) generally tolerated (F_N_ >0.1 and <10) or well enriched (F_N_ ≥ 10) substitutions on *E. coli* BW25113*ΔrfaD* and BW25113*ΔrfaG*. The upper p-value is the result of a Kruskal-Wallis test among all three groups while pairwise p-values from a Wilcoxon test are shown linking each group; p-values are shown if only if <0.05.

**Table S1.** Deep sequencing summary for phage variant expression library with and without DNAse treatment. Related to Figure 1, 2, and 4. Total number of reads passing filter, wild type reads, percentage wild type, total number of T7 acceptor reads and percentage of T7 acceptor reads in the T7 variant expression library with and without DNAse treatment (post ORACLE). See Table S3 for percentage distribution of each variant used in functional score calculations.

| Variant Expression Library |  | No DNase | DNase |
| --- | --- | --- | --- |
|  | Reads Passing Filter | 482152 | 298331 |
|  | Wild type Reads | 18481 | 10270 |
|  | Wild type % | 3.8% | 3.4% |
|  | Acceptor Reads | 5140 | 2641 |
|  | Acceptor % | 1.1% | 0.9% |

**Table S2.** Deep sequencing summary for phage variant library after selection under different conditions. Related to Figure 2 and 4. After selection on each host (shown left), for each biological replicate (R1, R2 and R3), total number of reads passing filter, wild type functional score and acceptor functional score. See Table S3 for F_N_ of each variant.

|  |  |  |  |  |
| --- | --- | --- | --- | --- |
| **10G** |  | **R1** | **R2** | **R3** |
|  | Reads Passing Filter | 275261 | 144033 | 239607 |
|  | Wild type functional score | 0.82 | 0.73 | 0.75 |
|  | Acceptor functional score | 0.000 | 0.005 | 0.007 |
| **BL21** |  | **R1** | **R2** | **R3** |
|  | Reads Passing Filter | 152679 | 476978 | 1067681 |
|  | Wild type functional score | 2.73 | 2.91 | 3.11 |
|  | Acceptor functional score | 0.004 | 0.008 | 0.009 |
| **BW25113** |  | **R1** | **R2** | **R3** |
|  | Reads Passing Filter | 180309 | 813142 | 687838 |
|  | Wild type functional score | 2.12 | 2.32 | 2.32 |
|  | Acceptor functional score | 0.009 | 0.011 | 0.008 |
| **BW25113ΔrfaG** |  | **R1** | **R2** | **R3** |
|  | Reads Passing Filter | 231007 | 347903 | 275610 |
|  | Wild type functional score | 0.12 | 0.07 | 0.09 |
|  | Acceptor functional score | 0.007 | 0.009 | 0.006 |
| **BW25113ΔrfaD** |  | **R1** | **R2** | **R3** |
|  | Reads Passing Filter | 330732 | 408389 | 260566 |
|  | Wild type functional score | 0.04 | 0.04 | 0.01 |
|  | Acceptor functional score | 0.007 | 0.006 | 0.007 |

**Table S3. Variant-specific details for phage variant expression library and variants after selection under different conditions. Related to Figure 1, 2 and 4**

See supplementary excel file

**Table S4. Functional comparison for each variant on different hosts and FD values. Related to Figure 3 and Figure 6**

See supplementary excel file

**Table S5. ΔΔG and FD Conversion for all variants. Related to Figure S5**

See supplementary excel file

**Table S6.**
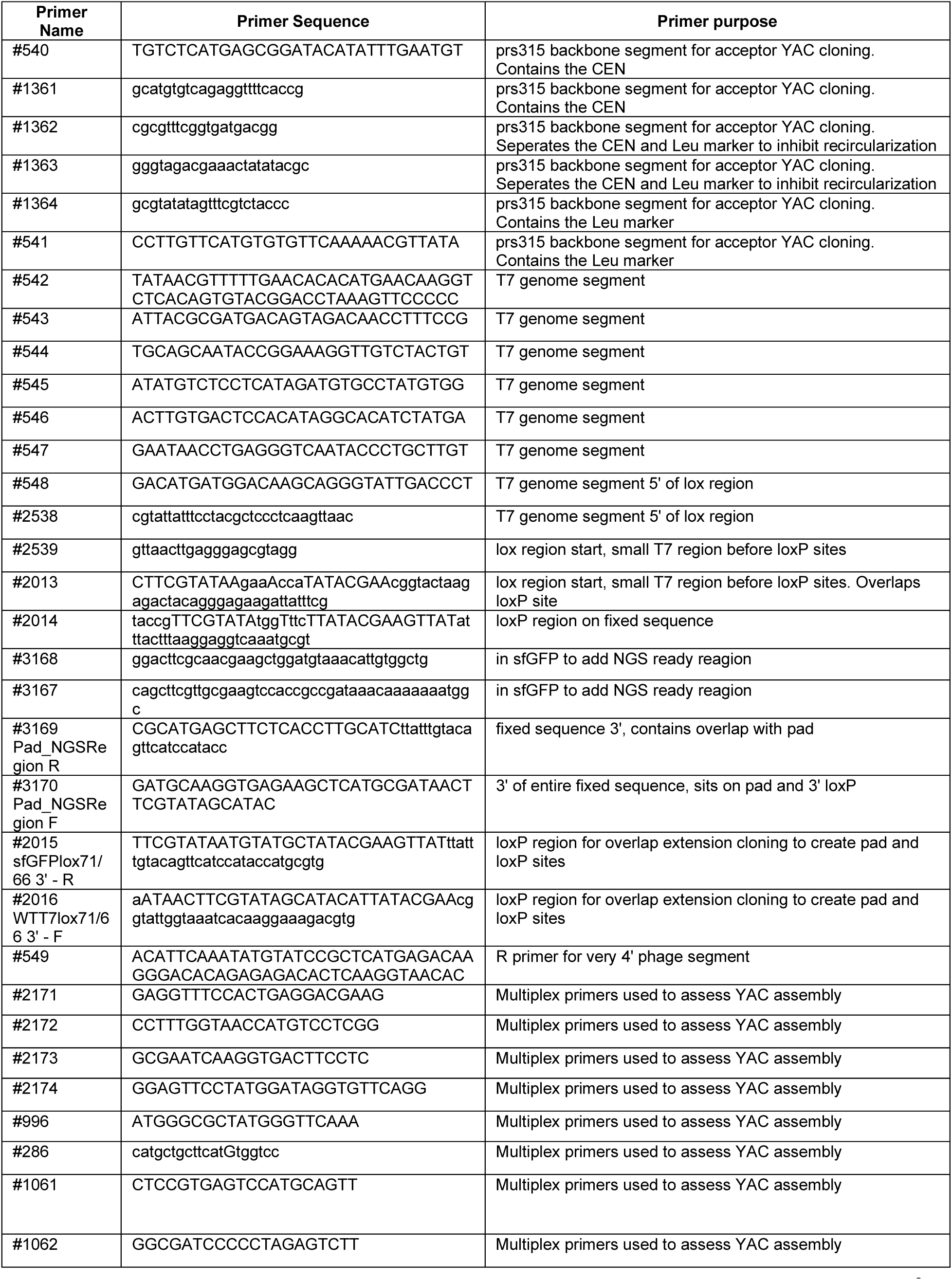
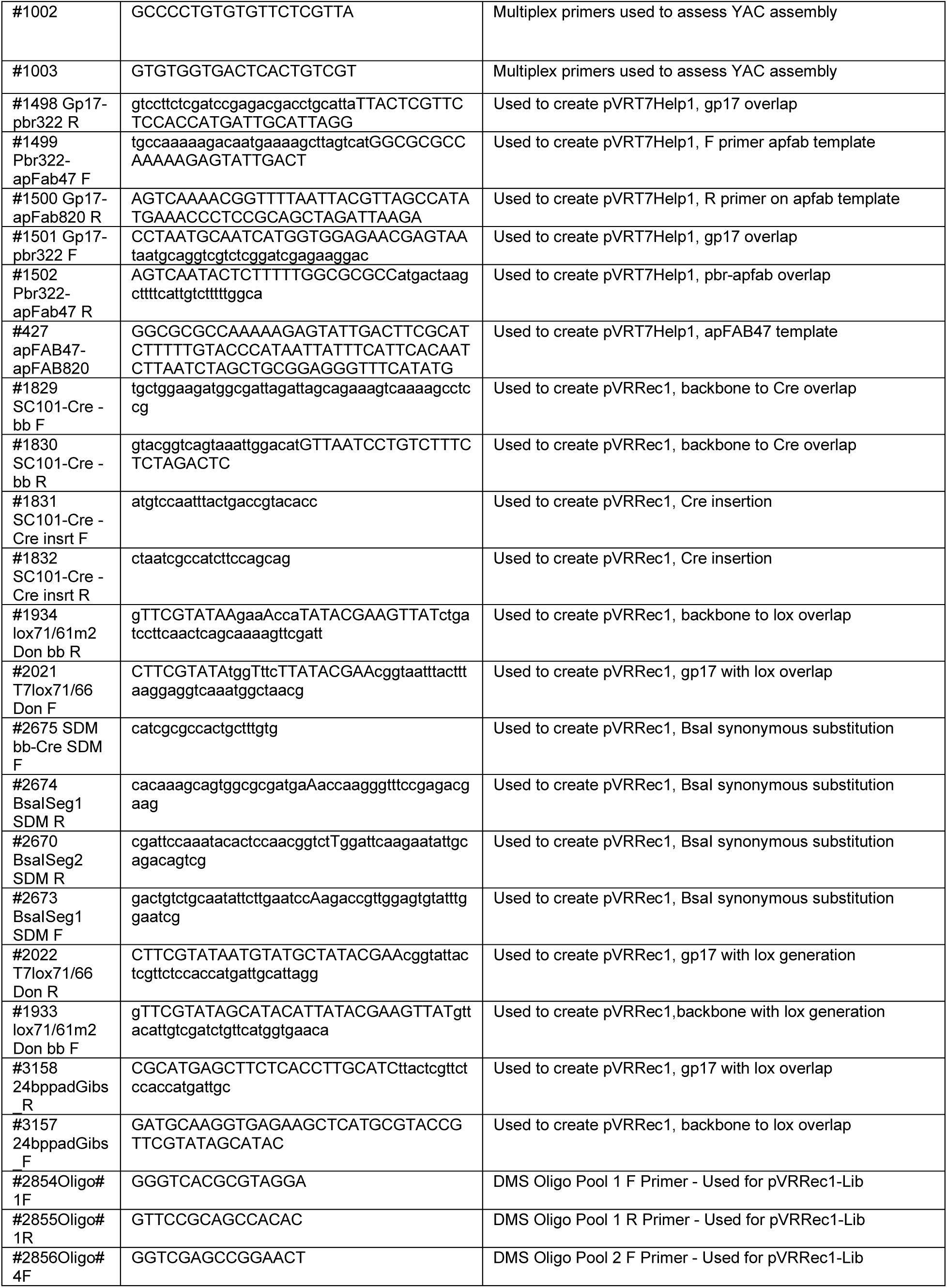
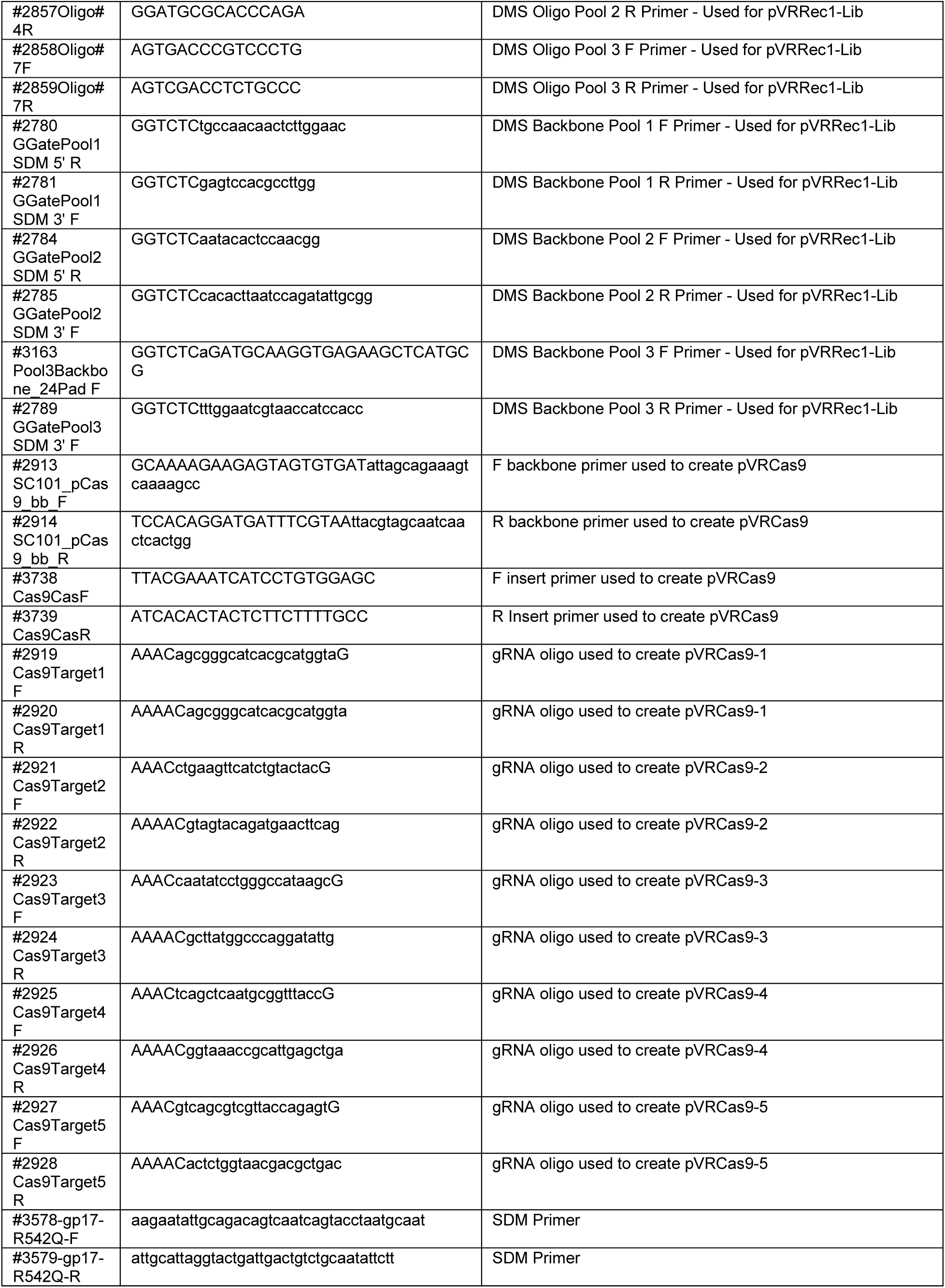
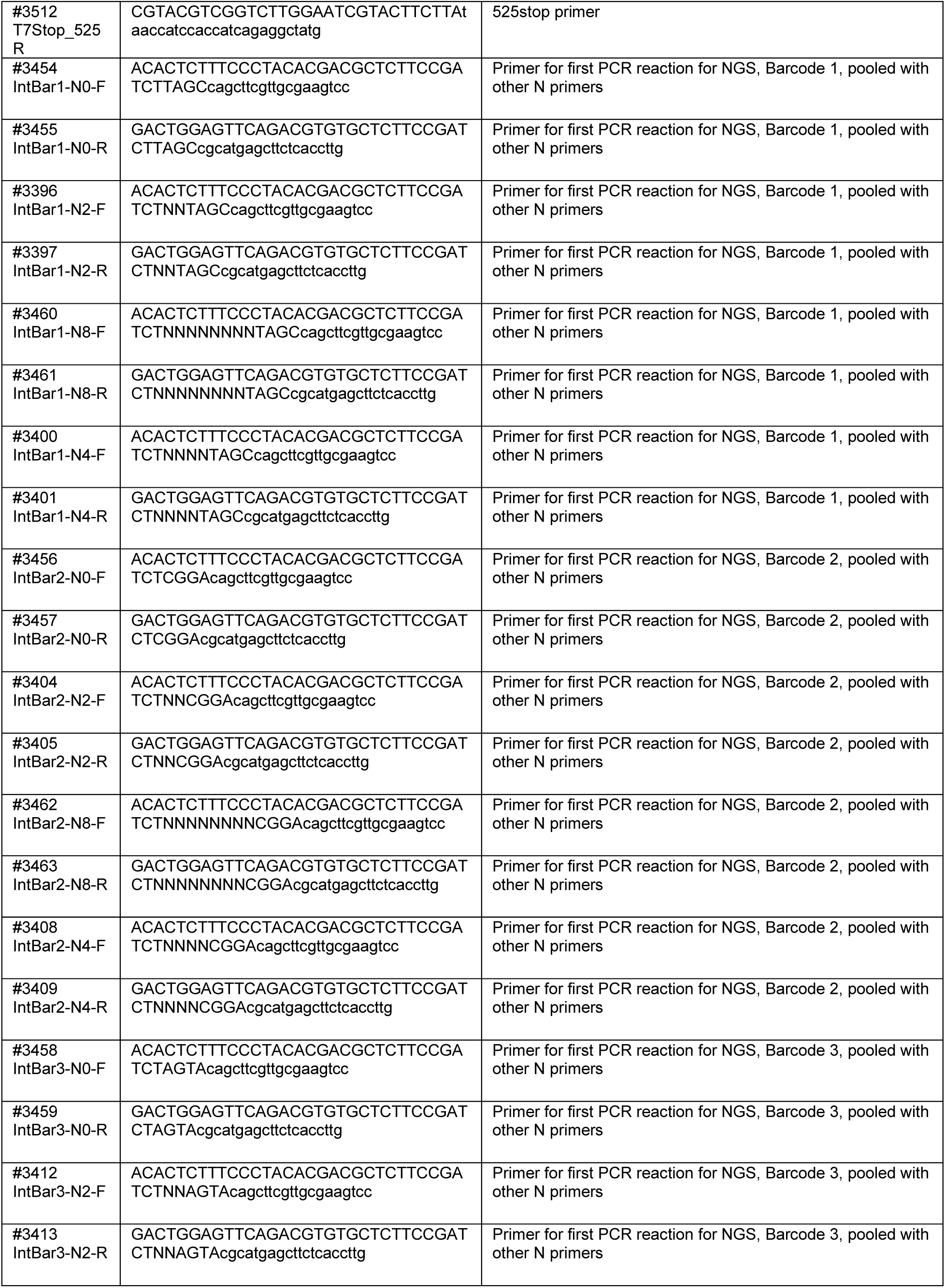
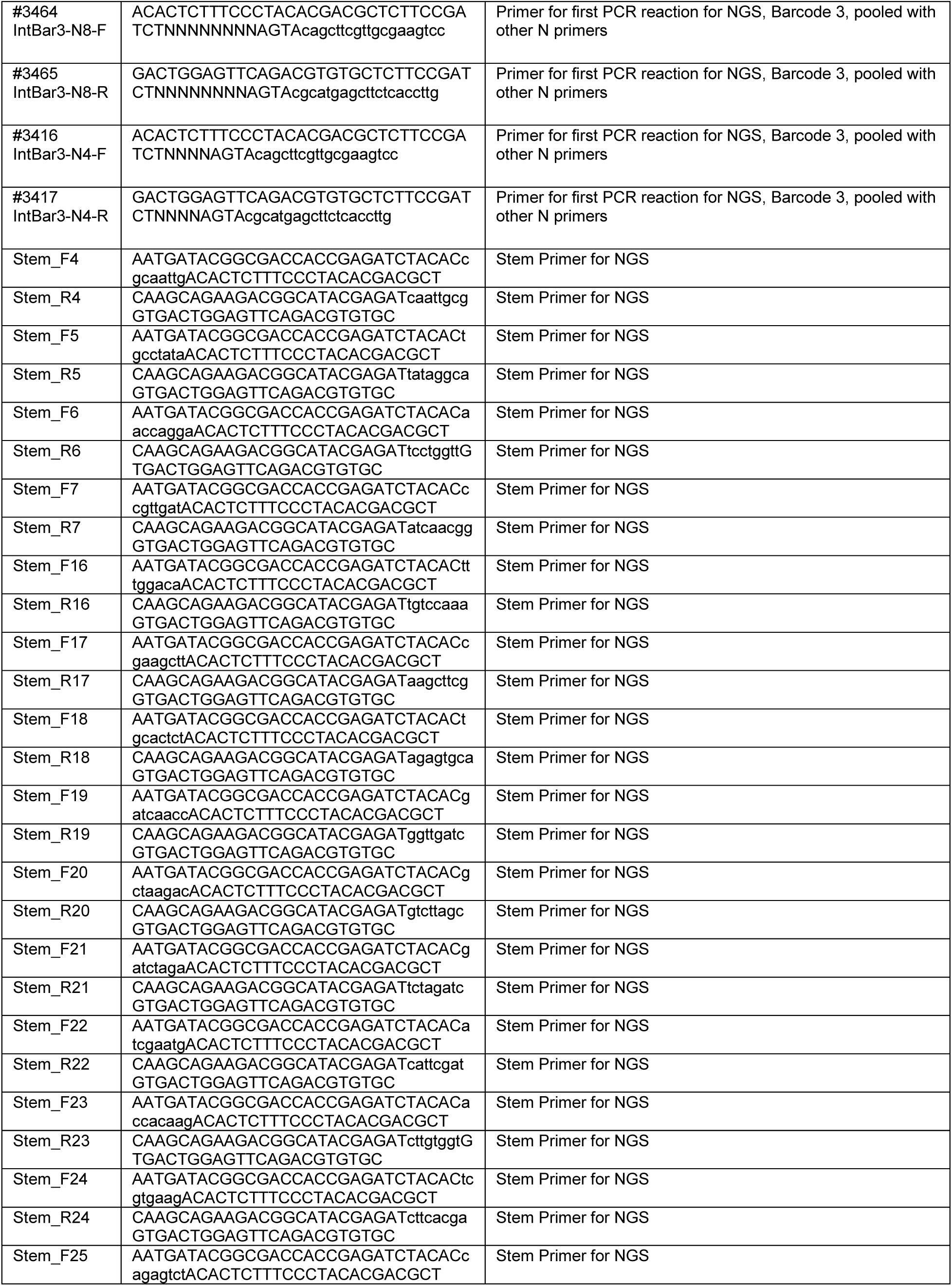
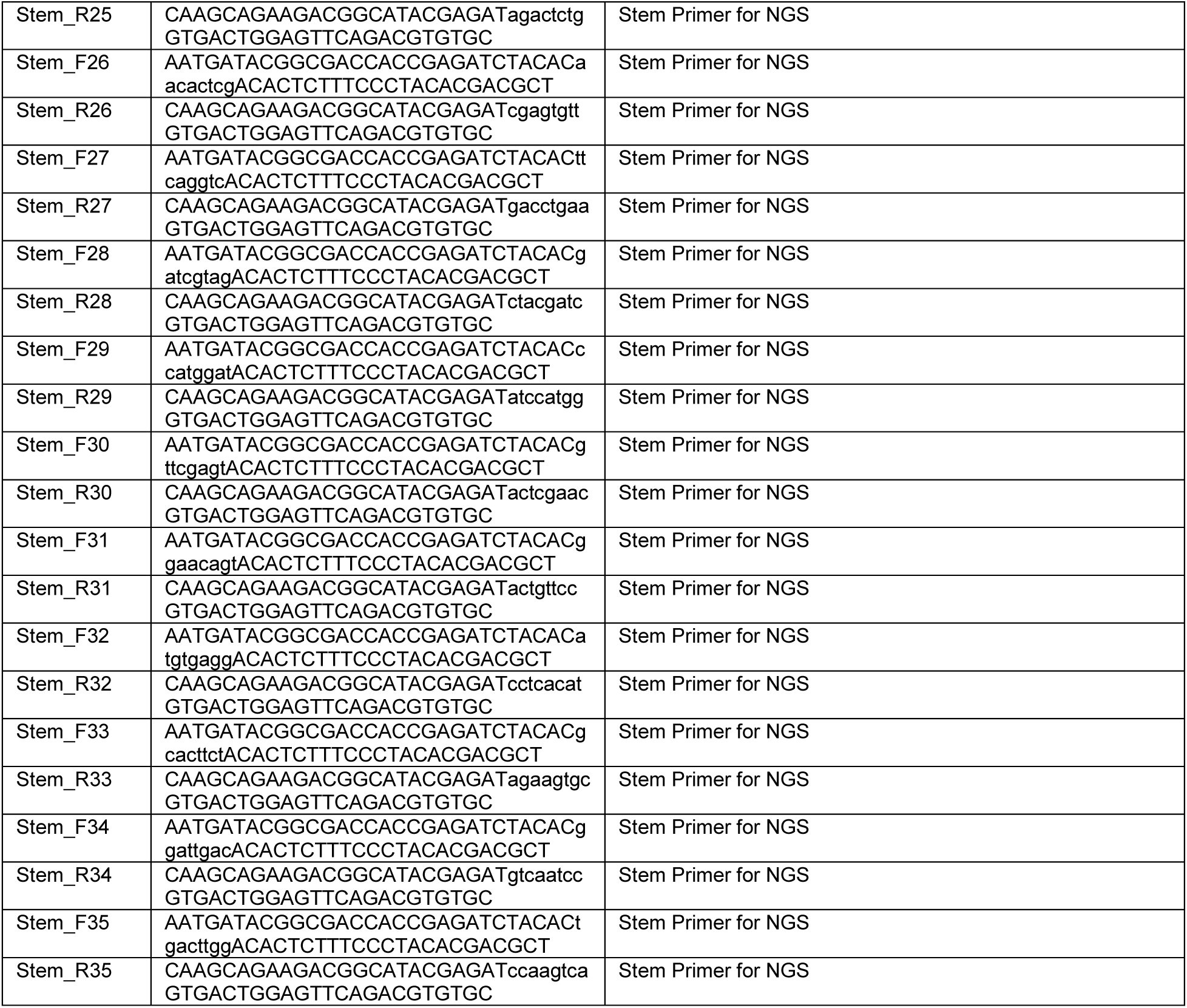
List of primers used for experimentation.

## Methods

### Microbes and Culture Conditions

T7 bacteriophage was obtained from ATCC (ATCC® BAA-1025-B2). *Saccharomyces cerevisiae* BY4741, *Escherichia coli* BL21 is a lab stock, *E. coli* 10G isa highly competent DH10B derivative [54] originally obtained from Lucigen (60107-1). *E. coli* 10-beta was purchased from NEB (C3020). *E. coli* BW25113, BW25113Δ*rfaD* and BW25113Δ*rfaG* were obtained from Doug Weibel (University of Wisconsin, Madison) and are derived from the Keio collection [55]. UTI473 was obtained from Rod Welch (University of Wisconsin, Madison) and originates from a UTI collection [50].

All bacterial hosts are grown in and plated on Lb media (1% Tryptone, 0.5% Yeast Extract, 1% NaCl in dH2O, plates additionally contain 1.5% agar, while top agar contains only 0.5% agar) and Lb media was used for all experimentation. Kanamycin (50 μg/ml final concentration, marker for pHT7Helper1) and spectinomycin (115 μg/ml final concentration, marker for pHRec1, pHRec1-Lib and pHCas9 and derivatives) was added as needed. All incubations of bacterial cultures were performed at 37°C, with liquid cultures shaking at 200-250 rpm unless otherwise specified. Bacterial hosts were streaked on appropriate Lb plates and stored at 4°C.

*S. cerevisiae* BY4741 was grown on YPD (2% peptone, 1% yeast extract, 2% glucose in dH2O, plates additionally contain 2.4% agar), after Yeast Artificial Chromosomes (YAC) transformation *S. cerevisiae* BY4741 was grown on SD-Leu (0.17% yeast nitrogen base, 0.5% ammonium sulfate, 0.162% amino acids – Leucine [Sigma Y1376], 2% glucose in dH20, plates additionally contain 2% agar). All incubations of *S. cerevisiae* were performed at 30°C, with liquid cultures shaking at 200-250 rpm. *S. cerevisiae* BY4741 was streaked on YPD or SD-Leu plates as appropriate and stored at 4°C.

T7 bacteriophage was propagated using *E. coli* BL21 after initial receipt from ATCC and then as described on various hosts in methods. All phage experiments were performing using Lb and culture conditions as described for bacterial hosts. Phages were stored in Lb at 4°C.

For long term storage all microbes were stored as liquid samples at -80°C in 10% glycerol, 90% relevant media.

SOC (2% tryptone, 0.5% yeast extract, 0.2% 5M NaCl, 0.25% 1M KCL, 1% 1M MgCl_2_, 1% 1M MgSO_4_, 2% 1M glucose in dH2O) was used to recover host and phages after transformation.

### General Cloning Methods

PCR was performed using KAPA HiFi (Roche KK2101) for all experiments with the exception of multiplex PCR for screening Yeast Artificial Chromosomes (YACs), which was performed using KAPA2G Robust PCR kits (Roche KK5005). Golden Gate assembly was performed using New England Biosciences (NEB) Golden Gate Assembly Kit (BsaI­HFv2, E1601L). Restriction enzymes were purchased from NEB with the exception of DNAse I (Roche 4716728001). DNA purification was performed using EZNA Cycle Pure Kits (Omega Bio-tek D6492-01) using the centrifugation protocol. YAC extraction was performed using YeaStar Genomic DNA Extraction kits (Zymo Research D2002). Gibson assembly was performed according to the Gibson Assembly Protocol (NEB E5510) but Gibson Assembly Master Mix was made in lab (final concentration 100 mM Tris-HCL pH 7.5, 20 mM MgCl_2_, 0.2 mM dATP, 0.2 mM dCTP, 0.2 mM dGTP, 10 mM dTT, 5% PEG-8000, 1 mM NAD^+^, 4 U/ml T5 exonuclease, 4 U/μl Taq DNA Ligase, 25 U/mL Phusion polymerase). All cloning was performed according to manufacturer documentation except where noted in methods. If instructions were variable and/or specific conditions are relevant for reproducing results, those conditions are also noted in the relevant methods section.

PCR reactions use 1 μl of ∼1 ng/μl plasmid or ∼0.1 ng/μl DNA fragment as template for relevant reactions. PCR reactions using phage as template use 1 μl of undiluted or 1:10 diluted phage stock, genomic extraction was unnecessary. Phage template was initially treated at 65°C for 10 minutes (for YAC cloning), but we later simply extended the 95°C denaturation step to 5 minutes (for deep sequencing).

DpnI digest was performed on all PCR that used plasmid as template. Digestion was performed directly on PCR product immediately before purification by combining 1-2 μl DpnI (20-40 units), 5 μl 10x CutSmart Buffer, PCR product, and dH2O to 50 ul, incubating at 37°C for 2 hours then heat inactivating at 80C for 20 minutes.

DNAse treatment of phages was performed by adding 5 μl undiluted phages, 2 μl 10x DNAse I buffer, 1 μl of 2 U/μl DNAse I, dH2O to 20 ul, then incubating for 20 minutes at 37°C, followed by heat inactivation at 75C for 10 minutes. 1 μl of this reaction was used as template for relevant PCR.

Electroporation of plasmids and YACs was performed using a Bio-rad MicroPulser (165-2100), Ec2 setting (2 mm cuvette, 2.5 kV, 1 pulse) using 25-50 μl competent cells and 1-2 μl DNA for transformation. Electroporated cells were immediately recovered with 950 μl SOC, then incubated at 37°C for 1 to 1.5 hours and plated or grown in relevant media.

*E. coli* 10G competent cells were made by adding 8 mL overnight 10G cells to 192 mL SOC (with antibiotics as necessary) and incubating at 21°C and 200 rpm until ∼OD_600_ of 0.4 as determined using an Agilent Cary 60 UV-Vis Spectrometer using manufacturer documentation (actual incubation time varies based on antibiotic, typically overnight). Cells are centrifuged at 4°C, 800g-1000g for 20 minutes, the supernatant is discarded, and cells are resuspended in 50 mL 10% glycerol. Centrifugation and washing are repeated three times, then cells are resuspended in a final volume of ∼1 mL 10% glycerol and are aliquoted and stored at -80°C. Cells are competent for plasmid and YACs.

Site Directed Mutagenesis (SDM) was performed, in brief, using complementary primers with the desired mutation in the middle of the primer, using 16x cycles of PCR, followed by DpnI digestion and electroporation into competent *E. coli* 10G. Splicing by Overlap Extension (SOE, also known as PCR overlap extension) was performed, in brief, using equimolar ratios of fragments, 16x cycles of PCR using extension based on the combined length of fragments, then a second PCR reaction using 1/100 or 1/1000 diluted product of the first reaction in a typical PCR reaction using 5’ and 3’ end primers for each fragment.

Detailed protocols for cloning are available on request. All primers used in experiments in this publication are listed in supplemental.

### Plasmid Cloning and Descriptions

<u>pHT7Helper1</u> contains a pBR backbone, kanamycin resistance cassette, mCherry, and the T7 tail fiber *gp17*. Both mCherry and *gp17* are under constitutive expression. *Gp17* was combined with promoter apFAB47 [56] using SOE and the plasmid assembled by Gibson assembly. There is a single nucleotide deletion in the promoter that has no effect on plaque recovery for phages that require *gp17* to plaque. This plasmid is used during optimized recombination and during accumulation in ORACLE to prevent library bias and depletion of variants that grow poorly on *E. coli* 10G.

<u>pHRec1</u> contains an SC101 backbone, Cre recombinase, a spectinomycin resistance cassette, and the T7 tail fiber *gp17* flanked by Cre lox66 sites with an m2 spacer, a 3’ pad region and lox71 sites with a wt spacer [53] (Figure S1). Cre recombinase is under constitutive expression. This plasmid was assembled with sequential PCR and Gibson assembly. During assembly we used PCR overhangs and SDM to create two synonymous substitutions in *gp17* to remove two BsaI restriction sites, facilitating downstream golden gate assembly. This plasmid was used in recombination assays, as it allows for recombination of wildtype *gp17*, and is used as template to generate the DMS variant library. The DMS variant library is referred to as pHRec1-Lib and is used during optimized recombination in ORACLE. Note this assembly was not tolerated in higher copy number plasmids.

<u>pHCas9</u> contains an SC101 backbone, a spectinomycin resistance cassette and cas9 cassette capable of ready BsaI cloning of gRNA [57]. This plasmid is used directly as part of the negative control for the accumulation assay, and has five derivatives, pHCas9-1 through -5, each with a different gRNA targeting the fixed region in the T7 acceptor phage. pHCas9 was created with Gibson assembly, while derivatives were assembled by phosphorylation and annealing gRNA oligos (100 uM forward and reverse oligo, 5 μl T4 Ligase buffer, 1 μl T4 PNK, to 50 μl dH2O, incubate at 37°C for 1 hour, 96C for 6 minutes then 0.1C/s temperature reduction to 23C), then Golden Gate cloning (1 μl annealed oligo, 75 ng pHCas9, 2 μl T4 DNA Ligase Buffer, 1 μl Golden Gate Enzyme Mix, dH20 to 20 ul. Incubation at 37°C for 1 hour then 60C for 5 minutes, followed by direct transformation of 1 ul, plated on Lb with spectinomycin). Note pHCas9-3 was the most inhibitory (Figure S2A) and was the only plasmid used in accumulation during ORACLE. This assembly was also not tolerated in higher copy number plasmids. All plasmid backbones and gene fragments are lab stocks.

### General Bacteria and Phage Methods

Bacterial concentrations were determined by serial dilution of bacterial culture (1:10 or 1:100 dilutions made to 1 mL in 1.5 microcentrifuge tubes in Lb) and subsequent plating and bead spreading of 100 μl of a countable dilution (targeting 50 colony forming units) on Lb plates. Plates were incubated overnight and counted the next morning. Typically, two to three dilution series were performed for each host to initially establish concentration at different OD_600_ and subsequent concentrations were confirmed with a single dilution series for later experiments.

Stationary phase cultures are created by growing bacteria overnight (totaling ∼20-30 hours of incubation) at 37°C. Cultures are briefly vortexed then used directly. Exponential phase culture consists of stationary culture diluted 1:20 in Lb then incubated at 37°C until an OD_600_ of ∼0.4-0.8 is reached (as determined using an Agilent Cary 60 UV-Vis Spectrometer using manufacturer documentation), typically taking 40 minutes to 1 hour and 20 minutes depending on the strain and antibiotic, after which cultures are briefly vortexed and used directly.

Phage lysate was purified by centrifuging phage lysate at 16g, then filtering supernatant through a 0.22 uM filter. Chloroform was not routinely used unless destruction of any remaining host was considered necessary and is mentioned in such cases.

To establish titer, phage samples were typically serially diluted (1:10 or 1:100 dilutions made to 1 mL in 1.5 microcentrifuge tubes) in Lb to a 10^-8^ dilution for preliminary titering by spot assay. Spot assays were performed by mixing 250 μl of relevant bacterial host in stationary phase with 3.5 mL of 0.5% top agar, briefly vortexing, then plating on Lb plates warmed to 37°C. After plates solidified (typically ∼5 minutes), 1.5 μl of each dilution of phage sample was spotted in series on the plate. Plates were incubated and checked at 2-4 hours and in some cases overnight (∼20-30 hours) to establish a preliminary titer. After a preliminary titer was established, phage samples were serially diluted in triplicate for efficiency of plating (EOP) assays. EOP assays were performed using whole plates instead of spot plates to avoid inaccurate interpretation of results due to spotting error [58]. To perform the whole plate EOP assay, 250 μl of bacterial host in stationary or exponential phase was mixed with between 5 to 50 μl of phages from a relevant dilution targeted to obtain 50 plaque forming units (PFUs) after overnight incubation. The phage and host mixture was briefly vortexed, briefly centrifuged, then added to 3.5 mL of 0.5% top agar, which was again briefly vortexed and immediately plated on Lb plates warmed to 37°C. After plates solidified (typically ∼5 minutes), plates were inverted and incubated overnight. PFUs were typically counted at 4-6 hours and after overnight incubation (∼20-30 hours) and the total overnight PFU count used to establish titer of the phage sample. PFU totals between 10 and 300 PFU were typically considered acceptable, otherwise plating was repeated for the same dilution series. This was repeated in triplicate for each phage sample on each relevant host to establish phage titer.

EOP was determined using a reference host, typically *E. coli* 10G with pHT7Helper1 but stated if otherwise. EOP values were generated for each of the three dilutions by taking the phage titer on the test host divided by the phage titer on the reference host, and this value was subsequently log_10_ transformed. Values are reported as mean ± SD.

MOI was calculated by dividing phage titer by bacterial concentration. MOI for the T7 variant library after the variant gene is expressed was estimated by titering on 10G with pHT7Helper1.

Limit of Detection (LOD) for T7 acceptor phages (T7 Acc) and T7 lacking a tail fiber (T7Δ*gp17*) was established based on the ability of these phages to clear a bacterial lawn. These phages are unable to plaque on host lacking pHT7Helper1, but phages do express a tail fiber due to being propagated on host with pHT7Helper1. Functionally this allows these phages to complete one infection cycle and kill one host but does not allow the creation of plaque-viable progeny phages if that host does not also contain pHT7Helper1. At an MOI of greater than ∼2 we noted plates no longer form lawns of bacteria but instead contain individual colonies or are clear, reflective of these singular assassinations. As expected, this effect occurs at different concentrations of phages for exponential or stationary host due to different host concentration at those stages of growth. As plaques cannot form under these conditions, and these infections are not productive beyond a single infection, we simply used this cut off as the limit of detection for this assay.

Growth time courses for UTI473 (Figure 5) and OD_600_ was performed using a Synergy HTX Multi-Mode 96-well plate reader, using 140 μl of host and 10 μl of relevant phage titers. Phages were applied after an hour of incubation in the plate reader.

### Recombination Rate and Accumulation Assays

To establish recombination rate (Figure 1C), we passaged T7 acceptor phages on 5 mL exponential phase *E. coli* 10G containing pHT7Helper1 and pHRec1. pHRec1 was used because the recombined *gp17* tail fiber is wildtype, ensuring every recombined phage is plaque-capable (derived from Figure S2B results). We sought to evaluate recombination rate after only 1 passage through the host to avoid misinterpretation of results in case recombined phages had different fitness than unrecombined phages. We used an MOI of 10 and allowed passage for 30 minutes, sufficient time for one wildtype phage passage, after which we halted any remaining reactions by adding 200 μl of chloroform and lysing the remaining bacterial host. Phages were then purified to acquire the final phage population. We established the phage population titer on 10G and 10G with pHT7Helper1. Both acceptor phages and recombined phages are capable of plaquing on 10G with pHT7Helper1 and this phage titer is used to count the total phage population. Only recombined phages are capable of plaquing on 10G and this titer is used to count recombined phages. Recombination rate was established as the fraction titer of recombined phages divided by recombined phages. This was repeated in triplicate and reported as mean ± SD.

> Method Note: It should be noted that this assay does not delineate for when recombination occurs in the host or how frequently recombination occurs in any one host. For example, if recombination were to occur on the original phage genome, all subsequent progeny phages could contain the recombined gene. In contrast, if recombination were to occur on the phage genome while it is being replicated, anywhere from one individual progeny to all progeny could contain the recombined gene.

To validate accumulation of recombined phages over acceptor phages (Figure 1D and S2E-F) we first generated a population of recombined phages using the same scheme as outlined for the recombination rate assay. After recombination this phage population contained primarily T7 acceptor phages with a small percentage of recombined phage containing a wildtype *gp17* tail fiber. This phage population was passaged on 10G containing pHT7Helper1 and either pHCas9-3 (targeting the fixed region in the acceptor phage using g3, the most effective guide by EOP, Figure S2A) or pHCas9 (randomized control). Phages were incubated with host in 5 mL total at an initial MOI of 1 based on the titer of the whole phage population. Every 30 minutes until 180 minutes, and thereafter every 60 minutes until 300 minutes, ∼250 μl of culture was removed, infection was stopped by adding 100 μl of chloroform, and phage samples were purified to establish the phage population at that timepoint. Titer at each timepoint was determined on both 10G and 10G with pHT7Helper1 with a single dilution series using whole plate plaque assay. Percent accumulation was derived by dividing titer on 10G by titer on 10G with pHT7Helper1. Accumulation on both hosts was repeated in triplicate and reported as mean ± SD.

### DMS Plasmid Library Preparation

To create the DMS variant plasmid library, oligos were first designed and ordered from Agilent as a SurePrint Oligonucleotide Library (Product G7220A, OLS 131-150mers). Every oligo contained a single substitution at a single position in the tip domain, overall including all non-synonymous substitutions, a single synonymous substitution, and a stop codon from position 472-554. Note we did mutate the stop codon, which is position 554, which when substituted results in a 3 amino acid extension (-DAR) of *gp17*. We used the most frequently found codon for each amino acid in the *gp17* tail fiber to define the codon for each substitution. Oligos contained BsaI sites at each end to facilitate Golden Gate cloning. To accommodate a shorter oligo length the library was split into three pools covering the whole tip domain. Oligo pools were amplified by PCR using 0.005 pmol total oligo pool as template and 15 total cycles to prevent PCR bias, then pools were purified. pHRec1 was used as template in a PCR reaction to create three backbones for each of the three pools. Backbones were treated with BsaI and Antartic Phosphatase as follows. 5 μl 10x CutSmart, 2 μl BsaI, ∼1177 ng backbone, dH2O to 50 μl was mixed and incubated at 37°C for 2 hours, after which 1 μl additional BsaI, 2 μl Antartic Phosphatase, 5.89 μl 10x Antartic Phosphatase buffer was spiked into reaction. Reaction was incubated for 1 more hour at 37°C, then enzymes were heat inactivated at 65°C for 20 minutes (concentration ∼20ng/μl at this point) and used directly (no purification) in Golden Gate Assembly. Golden gate assembly was performed using ∼100 ng of relevant pool backbone and a 2x molar ratio for oligos (∼10 ng), combined with 2 μl 10x T4 DNA ligase buffer, 1 μl NEB Golden Gate Enzyme mix and dH2O to 20 ul. These reactions were cycled from 37°C to 16°C over 5 minutes, 30x, then held at 60°C for 5 minutes to complete Golden Gate assembly. Membrane drop dialysis was then performed on each library pool for 75 minutes to enhance transformation efficiency. 2 μl of each pool was transformed into 33 μl competent *E. coli* 10-beta (NEB C3020) cells. Drop plates were made at this point (spotting 2.5 μl of dilutions of each library on Lb plates with spectinomycin) and total actual transformed cells were estimated at ∼2×10^5^ CFU/mL. Each 1 mL pool was added to 4 mL Lb with spectinomycin and incubated overnight, then plasmids were purified. Plasmids concentration was determined by nanodrop and pools were then combined at an equimolar ratio to create the final phage variant pool, denoted as pHRec1-Lib. pHRec1-Lib was transformed into *E. coli* 10G with pHT7Helper1. Drop plates were made (spotting 2.5 μl of dilutions of each library onto Lb plates with spectinomycin and kanamycin) and total actual transformed cells were also estimated at ∼2×10^5^ CFU/mL. The 1 mL library was added to 4 mL Lb with spectinomycin and kanamycin and incubated overnight. This host, *E. coli* 10G with pHT7Helper1 and pHRec1-Lib, was the host used for Optimized Recombination during ORACLE.

### ORACLE -Engineering T7 Acceptor Phages

Acceptor phages were assembled using YAC rebooting [17], [59], which requires yeast transformation of relevant DNA segments, created as follows. A prs415 yeast centromere plasmid was split into three segments by PCR, separating the centromere and leucine selection marker, which partially limits recircularization and improved assembly efficiency [60]. Wildtype T7 segments were made by PCR using wildtype T7 as template. At the site of recombination the acceptor phage contains, in order, lox71 sites with an m2 spacer [53] to facilitate one way recombinase mediated cassette exchange (RMCE), a fixed sequence that was derived from sfGFP with a nonsense mutation, a short region mimicking *gp17* to allow detection of acceptor phages by deep sequencing (5’ NGS in Figure S1), a 3’ ‘pad’ to facilitate deep sequencing, and lox66 sites with a wt spacer (see Figure S1). This entire region was turned into one DNA segment by serial SOE reactions.

> Method Note: reversable or two way RMCE using wildtype loxP sites could feasibly increase recombination efficiency, as one way RMCE is not necessarily required for ORACLE.

> Method Note: PCR, including PCR for deep sequencing, behaved inconsistently at lox sites. We theorize this may be because these sites are inverted repeats. Our inclusion of the 3’ ‘pad’ region corrected this problem and facilitated acceptor phage detection by deep sequencing.

DNA parts were combined together (0.1 pmol/segment) and transformed into *S. cerevisiae* BY4741 using the a high efficiency yeast transformation protocol [61] using SD-Leu selection. After 2-3 days colonies were picked and directly assayed by multiplex colony PCR to assay assembly. Multiplex PCR interrogated junctions in the YAC construct and was an effective way of distinguishing correctly assembled YACs. Correctly assembled YACs were purified and transformed into *E. coli* 10G cells containing pHT7Helper1, and after recovery 400 μl was used to inoculate 4.6 mL Lb. This culture was incubated until lysis, after which phages were purified to create the acceptor phage stock.

### ORACLE -Optimized Recombination

Recombination was performed by adding T7 Acceptor phages (MOI ∼5) to 15 mL exponential phase 10G with pHT7Helper1 and pHRec1-Lib (shown as the donor plasmid in Figure 1), split across three 5 mL cultures. A high MOI is used to allow for one effective infection cycle. Cultures were incubated until lysis (∼30 minutes). Lysed cultures were combined and purified. This phage population constitutes the initial recombined phage population and contained an estimated 2×10^7^ variants/mL in a total phage population of ∼2×10^10^ PFU/mL. The remainder of the phages are acceptor phages. A schematic of the recombination is shown in Figure S1. Note pHT7Helper1 ensures progeny should remain viable by providing *gp17 in trans*.

### ORACLE – Accumulation

Accumulation was performed by adding ∼MOI of 0.2 of recombined phages (50 μl or ∼1×10^9^ total phages) to 5 mL of stationary phase *E. coli* 10G with pHT7Helper1 and pHCas9-3 resuspended in fresh Lb with kanamycin and spectinomycin. Cultures are incubated until lysis (∼3.5 hours), then phages are purified. This MOI was chosen to target 1% of acceptor phages remaining in the final population as an internal control – the remainder of the phage population is accumulated variant phages. Stationary phase was used because it was more inhibitory based on EOP (Figure S2A). Note pHT7Helper1 still ensures progeny should remain viable by providing *gp17 in trans* and progeny from accumulation do not fully express variant genes.

> Methods Note: During ORACLE, the library gene is not actively repressed and some fraction of progeny phages are likely assembled with the variant gene, or contain chimeric tail fibers with both proteins. This may account for some degree of the skew we see during accumulation, although skewed residues were not consistent with enrichment patterns on 10G. Here, this skew was not significant, but this could be an influencing factor in future studies. Repression of the genomic library gene or decreasing accumulation time may be a viable option to enhance this protocol.

> Method Note: See Figure S2A for inhibition results for versions of pHCas9. When sequenced, individual plaques after selection had mutations in the region targeted by each gRNA, as expected for how resistance to Cas9 occurs. Note acceptor phages are not actively destroyed but are rather inhibited and maintained at the same concentration. Selection may be improved by using multiple guide RNAs or using sgRNA platforms.

### ORACLE -Library Expression

Library expression was performed by adding the accumulated DMS library to 5 mL *E. coli* 10G (with no plasmid) at an MOI of ∼1. Cultures are incubated for 30 minutes, then 200 μl chloroform is added to the culture to lyse any remaining cells and phages are purified. This constitutes the final phage variant library with full expression of the variant *gp17* tail fibers. This phage population is directly sequenced to establish the pre-selection library population.

> Method Note: This MOI and culture conditions are chosen to prevent phages from undergoing more than 1 replication cycle. Additional replication cycles would result in skew towards variants that grow better on the host used for expression. At an MOI of 1, progeny from the first replication cycle already comprise the majority of the possible concentration of phages for T7. Chloroform is added at 30 minutes to halt any rogue second infections in process.

> Method Note: Two points bear additional mention regarding ORACLE as a whole to create variant libraries. First, the importance of retaining variants that do not grow well on the host used to create the library cannot be overstated. These variants are critical for mapping functional regions. For example, we used 10G to grow our library, which happened to have the most significant selection of the susceptible hosts and had depletion of many functional regions. The resolution of this assay would have been deeply impacting if these variants had been lost. Second, due to possible depletion and skew, it is critical to assay library distribution after insertion and expression of the variants in the phage, instead of, for example, assaying distribution in plasmid before phage insertion. While ORACLE is designed to avoid this problem, any selection that occurs during library construction needs to be identified prior to selection experiments.

### DMS Selection

DMS selection was performed for all bacterial hosts in the same way. The T7 variant library was added to 5 mL of exponential host at an MOI of ∼10^-2^ and the culture was allowed to fully lyse (typically 40 to 80 minutes depending on the host). Phage lysate was purified and then the titer established for the host the phage was being selected on. This process was then repeated using the selected phage lysate. An MOI of 10^-2^ was chosen to allow phages which grow slower a chance to replicate. For reference under these conditions we expect wildtype to complete four infection cycles on a susceptible host. Phage lysate from the second selection was retained and used as template for deep sequencing to establish the post-selection phage population. The entire process was repeated in biological triplicate for each host.

### Deep Sequencing Preparation and Analysis

We used deep sequencing to evaluate phage populations. We first amplified the tip domain by two step PCR, or tailed amplicon sequencing, using KAPA HiFi. Primers for deep sequencing attach to constant regions adjacent to the tip domain (the target region is 304 bp total, between the 5’ NGS region and 3’ pad on Figure S1). Constant regions are also installed in the fixed region of the acceptor phages for the same size amplicon so acceptor phages can also be detected. The first PCR reaction adds an internal barcode (used for technical replicates to assay PCR skew), a variable N region (to assist with nucleotide diversity during deep sequencing, this is essential for DMS libraries due to low nucleotide diversity at each position), and the universal Illumina adapter. Undiluted phages are used as template. Four forward and four reverse primers were used in each reaction, each with a variable N count (0, 2, 4, or 8). Primers were mixed at equimolar ratios and total primers used was per recommended primer concentration. PCR was performed using 12 total cycles in the first PCR reaction, then the product of this reaction was then purified. The second PCR reaction adds an index and the Illumina ‘stem’. 1 μl of purified product from the first reaction was used as template using 8 total PCR cycles. The product of this reaction was purified and was used directly for deep sequencing. Each phage population was sampled at least twice using separate internal barcodes, and no PCR reactions were pooled. Total PCR cycles overall for each sample was kept at 20x to avoid PCR skew. All phage samples were deep sequenced using an Illumina Miseq System, 2×250 read length using MiSeq Reagent Kit v2 or v2 Nano according to manufacturer documentation.

Paired-end Illumina sequencing reads were merged with FLASH (Fast Length Adjustment of SHort reads) using the default software parameters [62]. Phred quality scores (Q scores) were used to compute the total number of expected errors (E) for each merged read and reads exceeding an Emax of 1 were removed [63]. Wildtype, acceptor phages, and each variant were then counted in the deep sequencing output. We correlated read counts for each technical replicate to determine if there was any notable skew from PCR or deep sequencing. Replicates correlated extremely well (R≥0.98 for all samples) indicating no relevant PCR skew. Besides wildtype and acceptor counts, we included only sequences with single substitutions in our analysis. While this limited the scope of the analysis, it greatly reduced the possibility of deep sequencing error resulting in an incorrect read count for a variant, as virtually every relevant error would result in at least a double substitution in our library. With this in mind, and to avoid missing low abundance members after selection, we simply used a low read cutoff of 2 and did not utilize a pseudo-count of 1 for each position.

Of the 1660 variants, three (S487P, L524M and R542N) fell below our limit of detection in the variant pool before selection. These positions were excluded from analysis as a pre-selection population could not be accurately determined, although both S487P and L542M subsequently emerged in several post-selection populations indicating they were present below the limit of detection. Technical replicates of each biological replicate were aggregated and each biological replicate was correlated to determine reproducibility (Figure S3). Lower correlation in 10G was noted to primarily be a result of differences in enrichment of several polar, uncharged substitutions and proline substitutions in the first biological replicate. These substitutions are proximally located near exterior loop HI and may indicate an unknown variable or growth condition in the first replicate that produces a slightly different enrichment pattern. Correlation was otherwise robust and excluding only G479Q produces R=0.94, R=0.95, and R=0.99 as displayed in Figure S3. Positively charged, downward facing substitutions were universally enriched in BW25113Δ*rfaD*, but correlation was reduced due to inconsistencies in which particular substitutions were the most enriched in a given replicate. While the same substitutions were enriched in all three replicates, suggesting reproducibility of results, the scale of enrichment varied considerably. Correlation was otherwise robust, excluding the single most enriched substitution of each replicate (G521R, A500R and G521H) produces R=0.90, R=0.93, and R=0.86 as displayed.

> Method Note: BW25113Δ*rfaD* is the most resistant host by EOP, and we hypothesize these inconsistencies in F_N_ may arise due severe loss of diversity, ‘bottlenecking’ the population and causing stochastic differences in enrichment to become magnified with multiple rounds of selection across independent experiments. Future work with very resistant hosts may benefit from fewer rounds of selection to prevent significant stochastic divergence.

To score enrichment for each variant we used a basic functional score (F), averaging results of the three biological replicates where 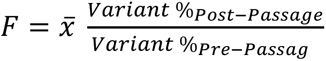. To compare variant performance across hosts we normalized functional score (F_N_) to wildtype, where 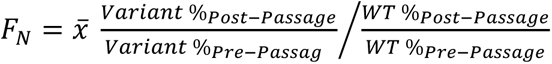.

### Classifying Variants and Isolating Variants

To define variant behavior on *E. coli* 10G, BL21, and BW25113 we considered variants depleted if F_N_ was below 0.1 (i.e. performed ten times worse than wildtype), tolerated if between 0.1 and 2, and enriched if above 2 (i.e. performed twice as good as wildtype) (Figure 2). As wildtype T7 effectively grows on all three hosts, we reasoned that it would be more challenging for an enriched mutant to surpass wildtype than it would be for a mutant to become depleted. These cutoffs were supported based on preliminary plaque assay results and the extent of standard deviation across biological replicates. For BW25113Δ*rfaD* and BW25113Δ*rfaG* we further defined significantly enriched variants as performing at least ten times better (F_N_≥10) than wildtype because wildtype does not grow effectively on these strains (Figure 4).

We compared variant F_N_ across 10G, BL21 and BW25113 to further characterize each variant and find functional variants (Figure 3). We sought to identify variants that had meaningfully different performance on different hosts, which would be strong evidence that either the wildtype residue or the variant substitution was important in a host-specific context. In addition to providing direct insight intro structure-function relationships, such substitutions or positions are ideal engineering targets for altering host range or increasing activity in engineered phages. We considered substitutions that were depleted (F_N_<0.1) on all three hosts to be intolerant, while substitutions that were tolerated or enriched (F_N_≥0.1) in all three hosts to be generally tolerated. Substitutions that were depleted on one host but tolerated or enriched on another were considered functional. To broadly characterize each position, we counted the number of substitutions at that position that fell into each category, and colored positions (Figure 3C) as functional if over 33% of substitutions at that position were functional, intolerant if over 50% of substitutions were intolerant, an tolerant otherwise. We found these cutoffs to effectively group residues of interest, although we note there remain substitutions that could be tolerant and relevant in different contexts or intolerant for these three hosts but not others.

For defining ideal host constriction mutants (Figure 6) we first constricted F_N_ values that were greater than 1 to reduce the impact of higher scores on this comparison. Specifically we generated Functional Difference (F_D_), where if F_N_ < 1, F_D_ = F_N_, and where 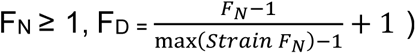. F_D_ thus ranged from 0 to 2 for each substitution where scores above 1 are normalized to the maximum value for that host and fall between 1 and 2, minimizing but not eliminating weight for enrichment. We reasoned that for the purposes of finding host constriction mutants, the extent of enrichment for a substitution is less relevant than if that substitution did poorly on another host. Put another way, it does not matter if a substitution is tolerated or enriched so long as it is depleted on a different host. For example, V544R has an F_N_ of 9.09 in *E. coli* 10G but 0.07 in *E. coli* BW25113, while G479E has an F_N_ of 1.73 in *E. coli* 10G and falls below the limit of detection for *E. coli* BW25113. For host constriction both positions should be scored highly, as the mutations can be tolerated or enriched in one host but are depleted in another. In contrast A500H has an F_N_ of 7.46 in *E. coli* 10G and 1.2 in *E. coli* BW25113. While F_N_ differs significantly and the substitution is enriched on one host, it is still tolerated in the other host and thus makes a poor host constriction target. After generating F_D_ we simply subtracted the substitution’s F_D_ on one host from the other in a pairwise comparison (Figure 6). Substitutions for host range constriction were considered ideal candidates if |F_D_| ≥ 1.

Variants with individual substitutions tested for EOP (Figures 4, 5 and 6) were either picked from plaques or created using SDM on pHRec1, which was used to create the variant using ORACLE.

### Rosetta ΔΔG and Physicochemical Property Comparison and Calculations

The crystal structure of the *gp17* tip domain was obtained (PDB ID: 4A0T) and water molecules removed before calculations run. All modeling calculations were performed using the Rosetta molecular modeling suite v3.9. Substitutions were generated using the standard ddg_monomer application [64] to enable local conformational to minimize energy. For comparison to F_D_ (Figure S5), a ΔΔG of 10 or greater was considered destabilizing and ΔΔG values between 10 and 30 were transformed to values between 0 and 1, with any ΔΔG greater than 30 set to 1. ΔΔG values below 10 were transformed to values between 0 and -1 based on a range to -9.29, the most stabilizing ΔΔG value calculated. We calculated F_D_ for this plot using the maximum F_D_ value from *E. coli* 10G, BL21 or BW25113. The maximum was used because any substitution that has a high F_D_ on any host was theorized to require a stable protein. Positions considered destabilizing (right side) are expected to have a very low maximum F_D_, whereas stabilizing positions (left side) may have a low or high F_D_ based on tolerance of the substitution. Positions that were tolerated or functional that Rosetta predicted to be unstable on the right of the plot may be due to errors in stability calculations or actual structural distortions that are either smaller perturbations that don’t affect fitness or are accommodated by quaternary arrangement. Alternatively, structural instability could be beneficial in some cases, allowing for enhanced receptor binding by, for example, exposing critical residues.

To compare physicochemical properties for *E. coli* 10G, BL21 and BW25113 we binned depleted (F_N_ ≤ 0.1) tolerated (F_N_ >0.1 and <2) or enriched (F_N_ ≥ 2) substitutions and derived the change in mass, hydrophilicity, and hydrophobicity for each substitution [65]. For BW25113Δ*rfaG* and BW25113Δ*rfaD* we binned using and F_N_ of 10 for the cutoff for enrichment instead. Packages ggplot2 and ggpubr in R were then used to compare the means of the three groups using a Kruskal-Wallis test [66], while subsequent pairwise comparisons were made using a Wilcoxon test [67].

### Statistical Analysis

Alluvial plots (Figure S4, Figure 4) and Figure 2F parallel plot were generated with RawGraphs. Violin plots for physicochemical properties are output from R. All other calculations and plots were made in Excel. Relevant statistical details of experiments can be found in the corresponding figure legends or relevant methods section.

